# Differences in firing efficiency, chromatin and transcription underlie the developmental plasticity of the Arabidopsis originome

**DOI:** 10.1101/258301

**Authors:** J. Sequeira-Mendes, Z. Vergara, R. Peiró, J. Morata, I. Aragüez, C. Costas, R. Mendez-Giraldez, J.M. Casacuberta, U. Bastolla, C. Gutierrez

## Abstract

Eukaryotic genome replication depends on thousands of DNA replication origins (ORIs) that constitute the originome. A major challenge is to learn ORI biology in multicellular organisms in the context of growing organs to understand their developmental plasticity. We have determined the originome and chromatin landscape of *Arabidopsis thaliana* at two stages of postembryonic development. ORIs associate with multiple chromatin signatures including TSS but also regulatory regions and heterochromatin, where ORIs colocalize with retrotransposons. In addition, quantitative analysis of ORI activity led us to conclude that strong ORIs have high GC content and clusters of GGN trinucleotides. Development primarily influences ORI firing strength rather than ORI location. ORIs that preferentially fire at early developmental stages colocalize with GC-rich heterochromatin whereas at later stages with transcribed genes, perhaps as a consequence of changes in chromatin features associated with developmental processes. Our study provides the originome of an organism at the postembryo stage that should allow us to study ORI biology in response to development, environment and mutations with a quantitative approach. In a wider scope, the computational strategies developed here can be transferred to other eukaryotic systems.

---

Replication of the large and complex genomes of multicellular organisms occurs during S-phase, once per cell cycle. Full genome replication is not achieved from a single DNA replication origin (ORI) but instead it depends on the coordinated function of thousands of ORIs scattered across the genome (Mechali 2010; Sanchez et al. 2012; Mechali et al. 2013). The association of pre-replication complexes (pre-RCs) with DNA determines the specification of genomic sites that can potentially act as ORIs. Some clues on the contribution of DNA sequence, chromatin marks, transcription factor binding sites and GC content to ORI specification and activity have been obtained (Mechali 2010; Costas et al. 2011; Leonard and Mechali 2013; Mechali et al. 2013; Gutierrez et al. 2016; Vergara and Gutierrez 2017). However, in spite of extensive efforts, the molecular determinants that specify the genomic location of ORIs in eukaryotes are still largely unknown. One of the reasons for this scarcity in our knowledge about ORI specification is the relative lack of well-characterized examples.

Genomic approaches carried out in various cultured cell models, including mammals, insects and plants, have helped to gain a picture supporting the view that ORIs frequently colocalize with chromatin marks associated with active chromatin (Sequeira-Mendes et al. 2009; Karnani et al. 2010; Macalpine et al. 2010; Cayrou et al. 2011; Costas et al. 2011; Cayrou et al. 2015; Comoglio et al. 2015). In addition, ORIs in the euchromatin of multicellular organisms are associated with GC-stretches, of which some may form G quadruplexes (Cayrou et al. 2012; Castillo Bosch et al. 2014; Valton et al. 2014). However there are large genomic regions where marks of active chromatin are absent but must be replicated, posing the question of whether there is a single chromatin signature able to define ORI location.

The use of cultured cells, where most studies have been done so far, to identify the mechanisms of ORI specification that function in the multicellular organism has some limitations (Mesner et al. 2013). Unlike *in vitro* culture conditions, cells within the body are subjected to hormonal signals and developmental cues that influence cell proliferation, cell fate decisions and differentiation. These intra- and extracellular factors, which are lost in the *in vitro* cell culture studies, can be crucial for the integration of genome replication with cell proliferation during development. Therefore, one major challenge is to identify ORIs in the cells of a whole organism to assess potential effects of cell fate acquisition and developmental cues on ORI specification.

Several studies have shown that in eukaryotes only a subset of ORIs is activated at each replication round. Thus, besides ORI specification, it is necessary to quantify the variability of ORI activity and investigate the factors that influence it. This has been recently approached in Caenorhabditis elegans by analyzing embryos of different ages (Pourkarimi et al. 2016; Rodriguez-Martinez et al. 2017) and in *Drosophila mealnogaster* polytene salivary glands (Sher et al. 2012). Here we have taken the challenge of defining the originome, that is the localization of all ORIs, and studying the variability of their activity in a living organism beyond the embryo stage. To identify ORIs in a whole organism and study their plasticity during postembryonic development we used the model plant *Arabidopsis thaliana* at two stages of vegetative development: 4 day-old seedlings (shortly after germination, when the hormonal and developmental signals necessary for vegetative growth have been established) and 10 day-old seedlings (before the transition to reproductive development). Since the young seedling contains a mixed population of dividing, endoreplicating, differentiating, embryo-derived and stem cells ((Gutierrez 2005); Fig. 1A), our approach should provide a collection of ORIs active in a wide variety of cell types and developmental stages. With this strategy we sought to obtain an understanding of (i) the molecular determinants of ORI specification and function, and (ii) their relationships with cell proliferation potential and the gene expression programs. We have found that ORIs are located in regions of all known chromatin states, indicating that there is not a single epigenetic signature that defines an ORI. Instead, different chromatin states do appear to affect the ORI relative strength because ORIs in active chromatin, though more prevalent, show lower signal intensity than the less abundant heterochromatin ORIs.

This suggests that once a location is chosen as an ORI in less accessible regions, it is maintained in more cells of the population. Finally, we have identified a subset of ORIs, preferentially active in 4 day-old seedlings, which are located in heterochromatin and another subset preferred in 10 day-old seedlings, associated with transcriptionally active genes. These results are supporting the association of ORI activity with certain DNA sequence, chromatin features and developmentally regulated transcriptional programs.

## Results

### Identification of ORIs and their replicative strength

Active ORIs are characterized by the presence of newly synthesized single-stranded DNA (ssDNA) molecules, also known as nascent strands (NS). NS purification from whole seedlings is challenging because of the limited amount of NS, even in highly proliferating cultured cells. Here, we have (i) implemented procedures to obtain sufficient amounts of a clean NS sample from whole plant seedlings, (ii) designed protocols to reduce possible biases associated with NS preparation and dsDNA conversion and (iii) developed computational tools to analyze ORIs in a quantitative manner.

NS were isolated from Arabidopsis seedlings in two stages of vegetative development: 4 day-old seedlings, soon after germination and 10 day-old seedlings, before the transition to reproductive development, which in both cases contain proliferating and endoreplicating cells of various differentiation stages and types. Our enhanced procedure yielded sufficient amounts of clean NS samples (see Methods, Fig. 1A and Supplemental Fig. S1). Briefly, after nuclei purification, NS were purified from DNA replication bubbles by sucrose gradient centrifugation to isolate DNA fragments of appropriate size in several gradient fractions (300 bp < nascent strands <2 kb, longer than Okazaki fragments but without compromising resolution). Any contaminating DNA fragmentation products were removed by *λ*-exonuclease (*λ*-exo) treatment because they are not protected by an RNA primer, as it occurs with *bona fide* NS. It has been reported that the *λ*-exo treatment produces a bias towards GC-rich DNA sequences (Foulk et al. 2015). However, this is significantly reduced provided that the treatment is carried out at least twice and under optimal conditions of substrate and X-exo concentrations (Cayrou et al. 2011; Picard et al. 2014; Cayrou et al. 2015; Comoglio et al. 2015; Lombrana et al. 2016).

**Figure 1.**
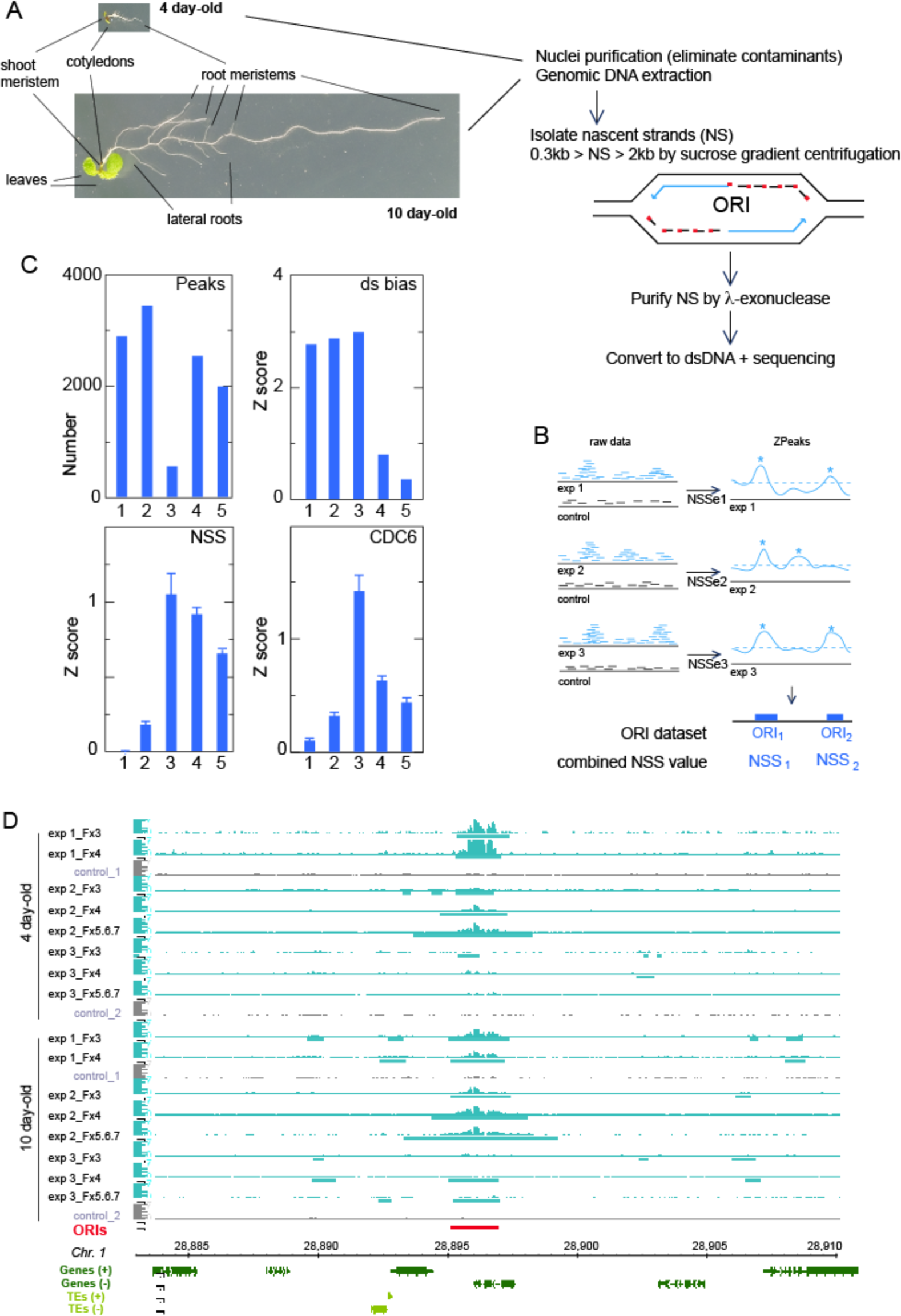
DNA replication origin (ORI) identification in whole Arabidopsis seedlings and evaluation of reproducibility and quality of sequencing datasets. (A) Summary of basic steps for purification of nascent strands (NS) from seedlings at two developmental stages of Arabidopsis vegetative growth. Seedlings contain cells undergoing cell proliferation and the endocycle in different locations. In 4 day-old seedlings, the shoot and the root apical meristems contain dividing cells whereas the cotyledons and the transition domain above the root apical meristem contain endocycling cells. The rest of the root is made up by different cell types some of them dividing and some differentiated. In addition to all these organs and cell types, 10 day-old seedlings contain growing leaf primordia and lateral root primordia, with proliferating cells, as well as a longer root with more differentiated cells. (B) Flowchart summary of the assignment of the nascent strand score (NSS) and the ZPeaks algorithm used to identify ORIs. (C) Quality controls of the NS purification and dsDNA conversion step. The horizontal axis represents the following datasets: class 1, biased peaks that do not overlap with the ORI set; class 2, all peaks with dsDNA conversion bias; class 3, biased peaks overlapping with ORIs; class 4, all ORIs; class 5, ORIs that do not overlap with the biased set. (D) Representative genome browser view of a ~35 kb region of chromosome 1, to illustrate ORI identification in various sucrose gradient fractions of the three independent experiments in 4 and 10 day-old seedlings.

Purified NS were further processed to generate libraries and submitted to sequencing Non-redundant sequence reads were aligned to the Arabidopsis genome (TAIR10) and those uniquely mapping were kept for further analysis. To allow a stringent identification of ORIs we carried out three independent experiments for each developmental stage and processed 2-3 consecutive sucrose gradient fractions in each case. In most cases, it was possible to detect the increase in the region covered by reads in fractions containing increased NS size, indicative of fork progression. Note that contaminating molecules derived from *λ*-exo-resistant non-nascent strands should have a similar region covered by reads. We reasoned that ORIs are in principle active in all the experiments, but with different probabilities of firing in the cell population. We identified a set of ORIs detected in at least two out of the three gradient fractions analyzed and then in, at least, two out three bological replicates (experiments) and quantified their strength in each experiment through the excess of NS reads with respect to the genomic control by calculating their NSS (Nascent Strand Score; Fig. 1B and Methods). In this way, the NSS values fully characterize all samples (two developmental stages and three independent experiments), whereas the locations of the ORIs do not vary since they are obtained by combining all samples. We computed weighted averages over the set of ORIs using the NSS as weights, so that ORIs that are weak in all samples contribute little to the weighted averages, thus reducing the chances of considering false positives. Conversely, the strong correlation that we found between NSS in different samples makes it unlikely that we missed any strong ORI, as we verified by visual inspection in the genome browser. To generate the originome we identified ORIs using our own peak-calling ZPeaks algorithm because (1) it provides a well-defined profile of the NSS over all of the genome, crucial for our analysis, and (2) it localizes an ORI at the local maximum of the NSS over the ORI box called, which is needed for carefully centering the metaplots (Fig. 1B).

We also carried out experiments to evaluate a possible bias in peak detection introduced during the dsDNA conversion step prior to library preparation. The routine protocol for dsDNA conversion reported in previous studies uses random primers mixed with heat-denatured genomic DNA and fast cooling to 37 °C. We found that this protocol has a bias to enrich for specific genomic regions when compared with the untreated genomic DNA and determined 6479 dsDNA-converted biased regions. Therefore, we searched for conditions that reduced the dsDNA conversion bias and found that carrying the primer-annealing step slowly (see Methods) halved the number of biased regions from 6479 to 3223. Therefore, we used this procedure with our NS samples, which we believe is advisable in all ORI mapping approaches that require a dsDNA conversion step. To remove any residual bias we used as background controls genomic DNA sheared, denatured and converted into dsDNA as for the NS samples. Importantly, the stringent strategy used to control for the dsDNA conversion bias has shown effective (Fig. 1C) since (i) ~80% of ORIs do not overlap with biased regions (class 5 in Fig. 1C) and possess high NSS and CDC6 (data from Costas et al., 2011) values, as expected for *bona fide* ORIs, (ii) likewise, ORIs that overlap with biased regions present even higher CDC6 binding and higher NSS, despite the dsDNA conversion bias contributes negatively to the NSS, and (iii) ~84% of biased regions do not overlap with ORIs (class 1 in Fig. 1C) and show a vanishing NSS score as expected for random genomic regions.

The ORI midpoint was identified with a resolution of 25 bp, the bin size used in the ZPeaks algorithm. We found that when an ORI was identified in different samples, the midpoint varied ~120 bp on average, which estimates the precision of our measurements. The ORI midpoint was then computed as the weighted average of the midpoints of the individual samples. The distribution of inter-ORI distance is highly skewed towards small values with a median of 27.6 kb much smaller than the mean of 43.4 kb. We tested ZPeaks both by visual inspection of the overlap between sequencing reads and candidate ORIs (Fig. 1D) and by statistical tests. Together our strategy allowed us to identify a robust set of 2374 bona fide ORIs that constitute the seedling originome (Supplemental Table 1) and can be confidently used to analyze the determinants of ORI specification.

We found that the NSS values are broadly distributed and span several orders of magnitude. Importantly, the NSS of different experiments are well correlated with each other, with correlation coefficients ranging from 0.52 to 0.71, with the exception of the comparison of experiments 2 and 3 using 4 day-old seedlings (Supplemental Fig. S2). This result lends support to our approach and suggests that some intrinsic ORI properties influence their firing rates in all the experiments although with important variations, as discussed below. Despite that the firing probability should saturate at 100% probability, the NSS did not show any sign of saturation, suggesting that they are far from 100% activity. To facilitate the comparison between the two developmental stages, we obtained scores that averaged the three independent experiments performed for each stage, as described in Methods.

### Validation of ORI activity

We wanted to validate ORI activity in an independently prepared sample of nascent strands by measuring the relative abundance of nascent strands by qPCR across genomic regions that contained ORIs identified by SNS-seq. This approach can be used for a reduced number of ORIs but has the advantage of using a nascent strand preparation that consists of ssDNA, treated with *λ*-Exo but that has not been converted into dsDNA. To assess ORI activity in a stringent way, we selected ORIs belonging to either the class of ORIs not overlapping with dsDNA-converted biased regions or to ORIs overlapping with those regions. As a control, we chose a region lacking any significant amount of reads. Nascent strands were prepared independently of those used for the SNS-seq experiments from 4 and 10 day-old seedlings and used as substrate in qPCR amplification reactions using primer pairs spanning ~10-15 kb around the ORI sites (Fig. 2 and Supplemental Table 2). Remarkably, we detected clear peaks corresponding to active ORIs, including ORIs that overlapped with dsDNA-converted biased regions. These data demonstrate that the originome generated under our stringent conditions constitutes a *bona fide* genome-wide set of ORIs active in Arabidopsis seedlings.

**Figure 2.**
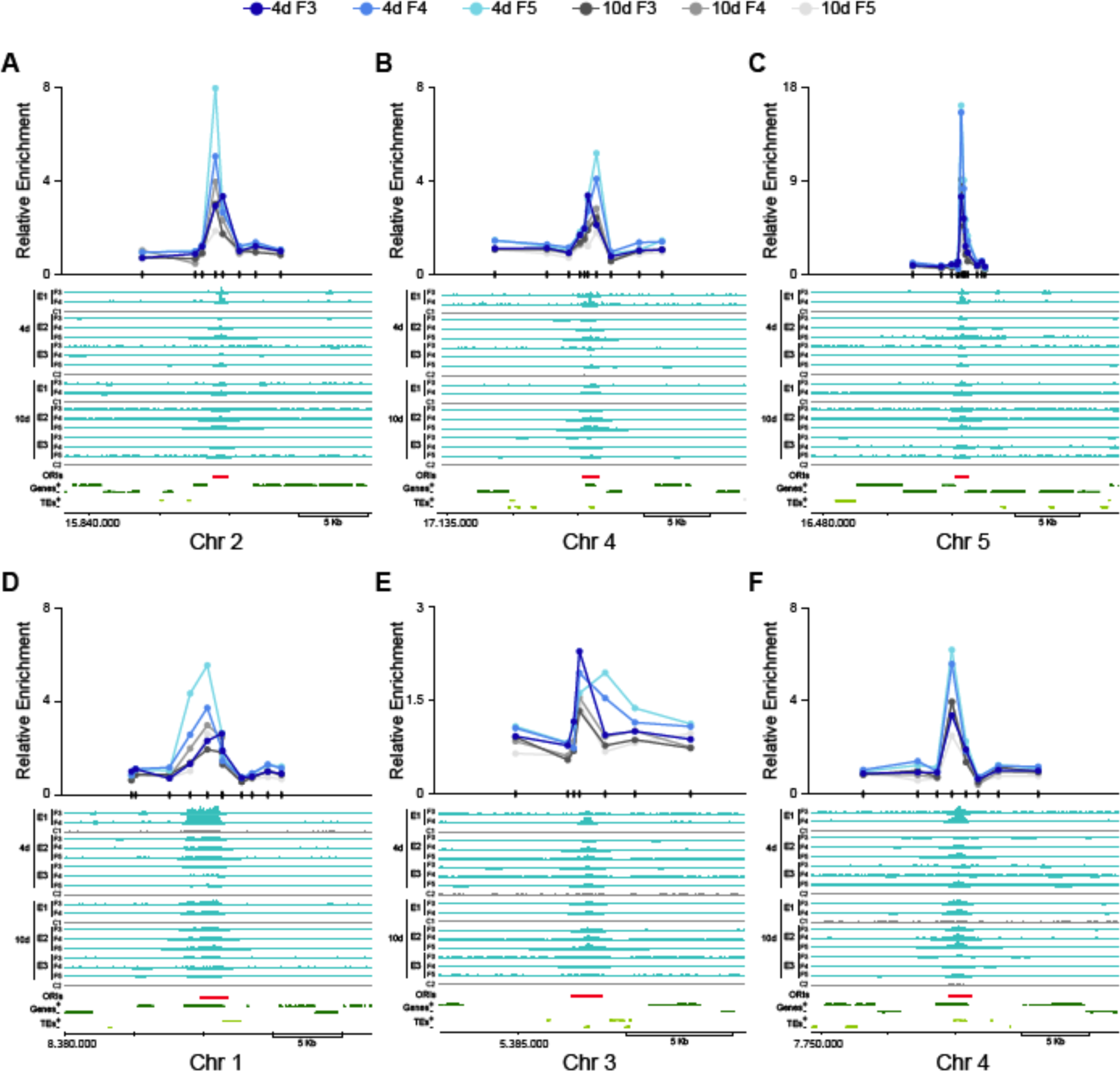
Validation of DNA replication origin activity by measuring nascent strands abundance. Several ORI-containing regions were chosen from the SNS-seq datasets belonging to ORIs (red bars at the bottom) that did not (A-C) coincide with biased regions (class 5 in Fig. 1C) and that colocalize with regions with some background in the controls (D-F; class 3 in Fig. 1C) to determine the abundance of nascent strands by qPCR. The sample was an independent preparation of nascent strands, purified by sucrose gradient centrifugation, treated with *λ*-exonuclease but not converted to dsDNA. The genomic regions under study are shown at the bottom of each panel and contain the read maps across them for the various sucrose gradient fractions used of the three independent experiments from 4 and 10 day-old seedlings, as indicated. Coordinates and location of primer pairs used to scan each region (small marks in the X-axis) are also indicated. A region lacking SNS-seq signal was used as a negative control and to normalize the qPCR values.

### Genomic landscape of ORI locations

To determine the preferences of ORI localizations, we examined the association of ORIs with various genomic elements. Most ORIs (>78%) are associated with genic regions, including 1 kb upstream regions, much more than expected by chance. Within genes, ORIs locate more frequently in exons (Fig. 3A). Intergenic regions and TEs comprise ~5% and ~13% of ORIs, respectively, while these genomic regions represent a much larger fraction of the genome (~15% and ~21%, respectively). We recently discovered that ~5% of ORIs active in cultured Arabidopsis cells colocalize with TEs, although in the gene-poor pericentromeric heterochromatin the frequency of colocalization with TEs increases to ~34%. Moreover, these ORIs are much more frequently located within retrotransposons than in DNA transposons, and among them in TEs of the Gypsy and LINE families (Vergara et al. 2017). Thus, we use the same approach to investigate the distribution of ORIs active in seedlings and found that in pericentromeric regions the frequency of ORIs located within TEs increases significantly compared to the overall genome (Fig. 3B). Furthermore ORI-TEs show a preference to colocalize with retrotransposons, as it occurs in cultured cells, although in seedlings the fraction of ORIs located in DNA transposons is higher, in particular for the MuDR family (Fig. 3C).

**Figure 3.**
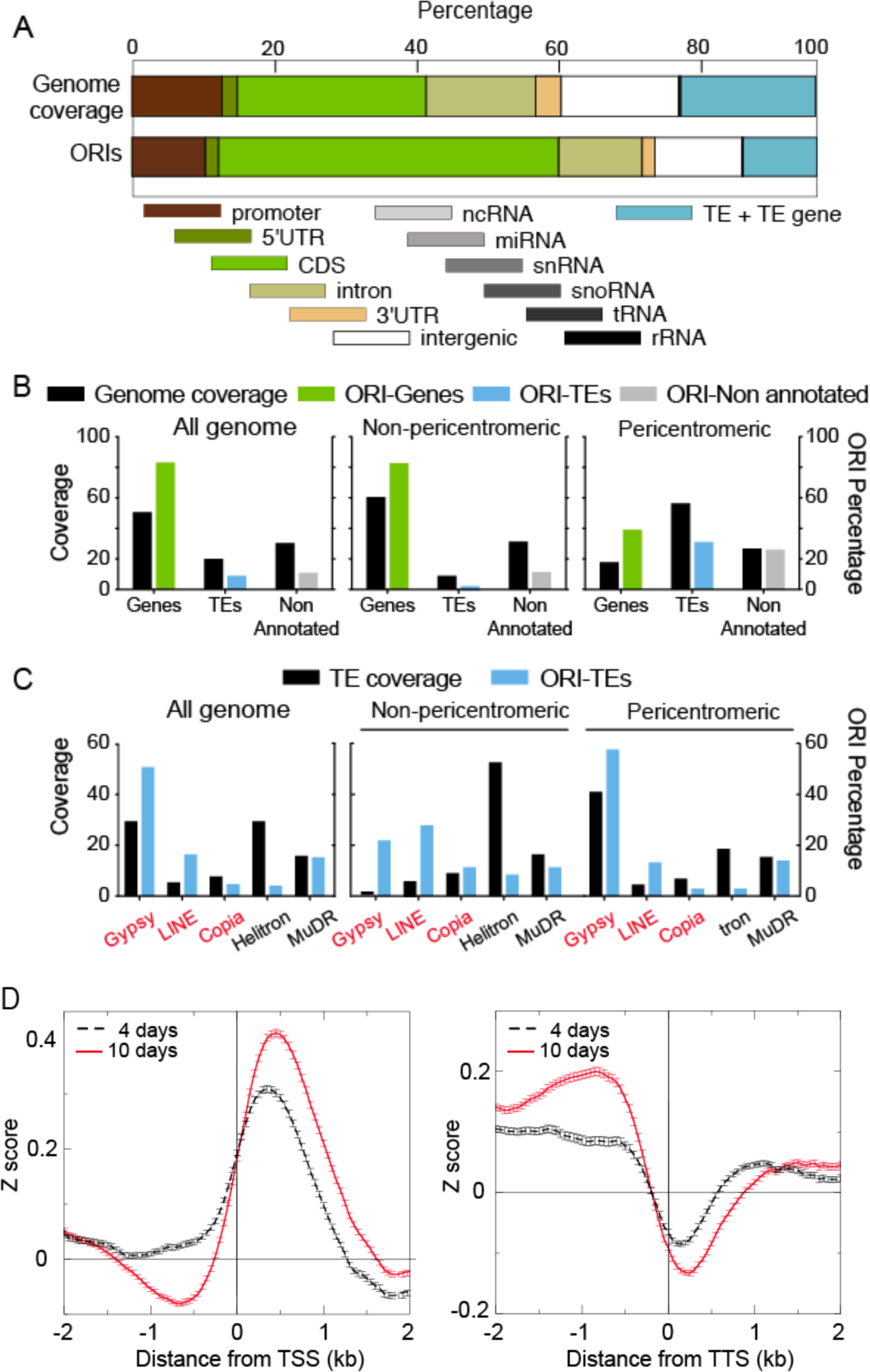
Association of ORIs with genomic elements and TE families. (A) Relationship between ORI location and genomic elements. The overlap (in base pairs) between the indicated genomic elements and each ORI was computed and expressed as a percentage. A region of 1 kb upstream the coding sequence was considered as the promoter. Note that TEs are large genomic elements that may have one or more TE genes associated with them. Here, the class TE refers to genomic regions that contain TEs but do not overlap with TE genes. (B) Frequency distribution of ORIs colocalizing with genes (green), TEs (blue) and non-annotated regions (grey) compared with the respective nucleotide coverage. (C) Frequency distribution of ORI-TEs (blue bars) in TE families in all the Arabidopsis genome, the non-pericentromeric regions and the pericentromeric regions compared with the respective TE family nucleotide coverage of total TE nucleotides (black bars). In the X-axis, retrotransposon families (red) and DNA transposon families (black). (D) Metaplots of the combined NSS of the three independent experiments of 4 day-old and 10 day-old seedlings with respect to the transcription start sites (TSS; left panel) or the transcription termination site (TTS; right panel), oriented in both cases with the transcribed RNAs.

We also found that ORIs have a preference to localize ~0.5 kb downstream from the Transcription Start Site (TSS) and tend to avoid Transcription Termination Sites (TTS) at both developmental stages (Fig. 3D).

### Local properties of ORI locations

To study the properties of Arabidopsis ORIs in seedlings, we assessed the average local neighborhood of all ORIs by computing metaplots centered at the ORI midpoint with each ORI being weighted with its own NSS. To account for the asymmetry between the G and C and between the A and T bases, known to be a feature of replication origins, ORIs were oriented along the 5’-3’ direction of the strand with positive GC skew. For this analysis each variable was transformed into a Z score with respect to the entire genome, which allows comparing the strength of different genomic and epigenomic variables.

First of all, we confirmed that the NSS of all experiments had a very prominent peak at the ORI midpoint (Fig. 4A and Supplemental Fig. S3). Additionally, we examined the position of the pre-RC protein CDC6 (Costas et al. 2011). As expected, we found that CDC6 has a high peak centered at the midpoint of active ORIs with a width of ±1 kb (Fig. 4A and Supplemental Fig. S3). This provides strong independent support to the peak-calling procedure used here to define ORI location based on nascent-strand mapping. We also found that the location of ORI midpoint coincides with a peak of nucleosome occupancy (Fig. 4A and Supplemental Fig. S3), as described for cultured cells in Arabidopsis and mammals (Stroud et al. 2012; Lombrana et al. 2013). Furthermore, the regions around ORIs are more frequently transcribed than the average of the genome, with a broad peak of approximately ±1 kb centered at the ORI midpoint (Fig. 4A and Supplemental Fig. S3).

**Figure 4.**
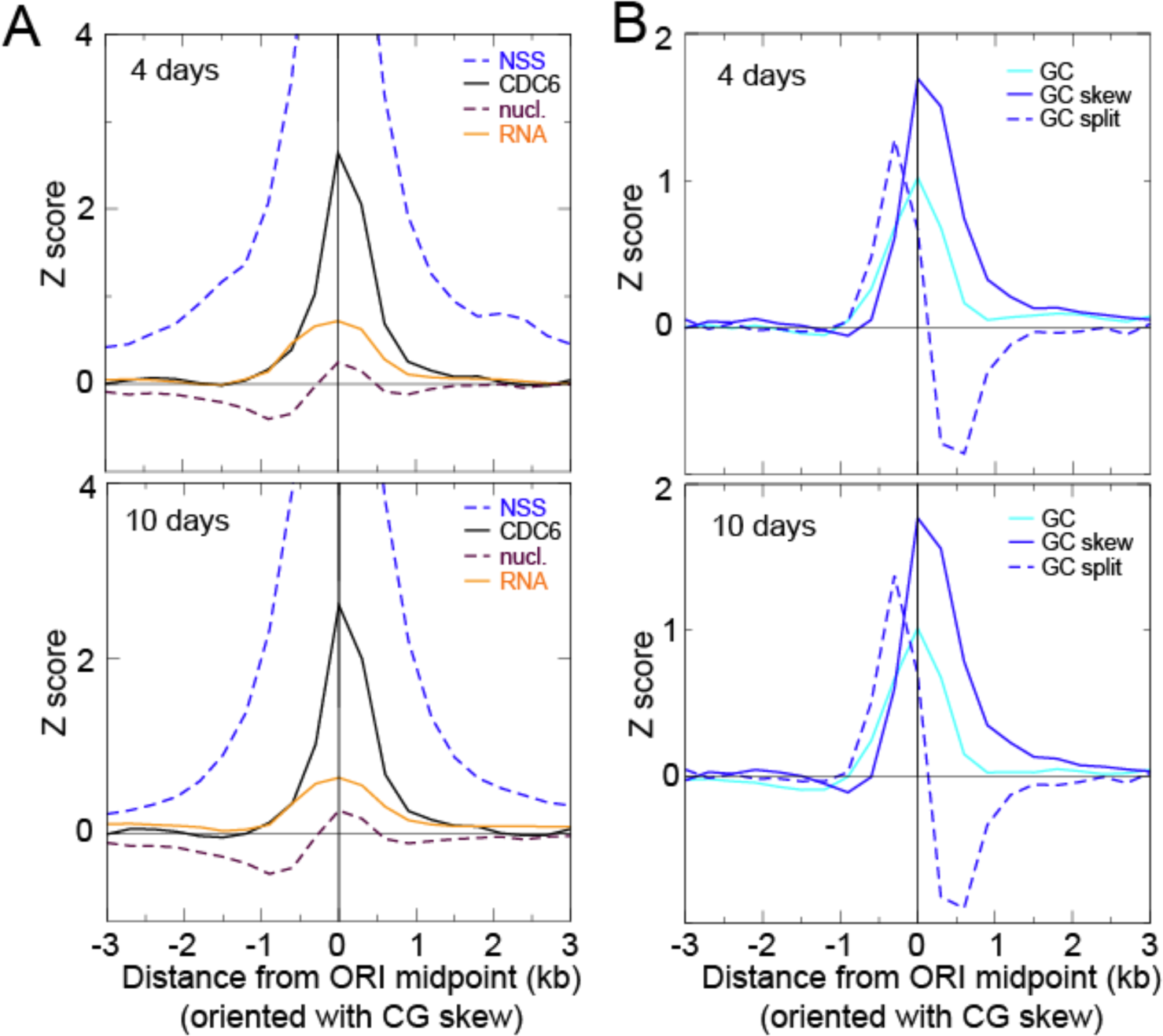
Features of the local neighborhood of ORIs in whole Arabidopsis seedlings. (A) Metaplots of NSS, CDC6, transcript content (RNA) and nucleosome (nucl.) content weighted with the combined Z score of the three independent experiments of 4 day-old (top) and 10 day-old (bottom) seedlings. The metaplots for individual scores of each experiment are shown in Supplemental Fig. S3. (B) Metaplots of GC, GC skew and GC split weighted with the combined Z score of the three independent experiments of 4 day-old (top) and 10 day-old (bottom) seedlings. The metaplots for individual scores of each experiment are shown in Supplemental Fig. S4.

The Arabidopsis genome is particularly rich in A-T (63.8%). ORIs identified in cultured cells preferentially colocalize with short G+C-rich stretches (Costas et al. 2011). Measuring the G+C content in 100 bp sliding windows across all ORI locations revealed that ORIs in seedlings also colocalize with G+C-rich regions that show a peak of ~0.8 kb in width, centered at the ORI midpoint (Fig. 4B and Supplemental Fig. S3). The asymmetry between the G and C nucleotide, called GC skew, is a signature of ORIs both in prokaryotes and metazoa (Macalpine et al. 2010; Cayrou et al. 2011; Arakawa and Tomita 2012; Xia 2012; Comoglio et al. 2015). The GC skew presents a strong peak centered at the ORI midpoint but asymmetrically distributed (−0.5 kb through 1.0 kb from ORI midpoint).

The GC skew associated with DNA replication is attributed to the asymmetry of mutation processes on the leading and lagging strand. To test this we defined the GC split as the difference of GC skew downstream and upstream of a genomic point. Under the hypothesis that the GC skew is produced by the different mutation processes in the leading and lagging strands, we expect that the GC split has a maximum coinciding with the ORI midpoint. However, we observed that ORI midpoints typically lay in between a strong maximum of the GC split upstream of the ORI midpoint at approximately −0.3 kb, and a slightly less strong minimum at ~0.4 kb (Fig. 4B and Supplemental Fig. S3). This result does not support the hypothesis that the different mutation processes at the leading and lagging strand are the sole cause of the GC skew.

### ORIs associate with multiple chromatin signatures

The mechanisms responsible for ORI specification along the genome in multicellular eukaryotes remain unknown. Studies in several model organisms, including mammals, Drosophila and plants, have revealed a preferential association of ORIs with activating chromatin marks (Sequeira-Mendes et al. 2009; Cayrou et al. 2011; Costas et al. 2011; Picard et al. 2014; Cayrou et al. 2015; Comoglio et al. 2015; Pourkarimi et al. 2016; Rodriguez-Martinez et al. 2017). A simplistic interpretation of these observations may suggest that some combination of chromatin features may be sufficient for ORI specification. However, simple inspection of genomic data clearly shows that there is not a single epigenetic mark or a simple combination of them common to all ORIs. Recent studies demonstrate the existence of three major classes of ORIs with different organization, chromatin environment, and sequence motifs (Cayrou et al. 2015), suggesting that ORIs are associated with different signatures.

To investigate the preferences of ORIs to occur in particular chromatin settings, we assigned each ORI midpoint to one of the chromatin states defined by high-resolution analysis of 16 chromatin and DNA features in the Arabidopsis genome (Sequeira-Mendes et al. 2014). These states simplify the combinatorial complexity of DNA and histone marks across the Arabidopsis genome into nine chromatin states characterized by unique signatures, in a manner similar to what has been done for Drosophila and human cells (Ernst et al. 2011; Kharchenko et al. 2011). Importantly, the chromatin states identified in Arabidopsis shows a preferred linear sequence defining proximal promoters (state 2) – TSS (state 1) – 5’end of genes (state 3) – long genes (state 7) – 3’end of genes (state 6), followed by repressed states containing Polycomb marks (states 5 and 4) and two types of heterochromatin (states 8 and 9; (Sequeira-Mendes and Gutierrez 2016)).

The cumulative weight of ORIs in the chromatin states is higher for ORIs colocalizing with state 1 (TSS), which are the most numerous (Fig. 5A and Supplemental Fig. S4A). Moreover, ORIs that colocalize with state 2 (proximal promoters and 5’UTRs) and state 3 (5’-end of genes) also have a relatively high weight (Fig. 5A and Supplemental Fig. S4A). On the contrary, state 5 (PcG-repressed regions) and states 8 and 9 (the two heterochromatin types), state 7 (long coding genes), state 6 (3’ end of genes) and in particular state 4 (distal regulatory intergenic regions) contain more moderate amounts of ORIs.

**Figure 5.**
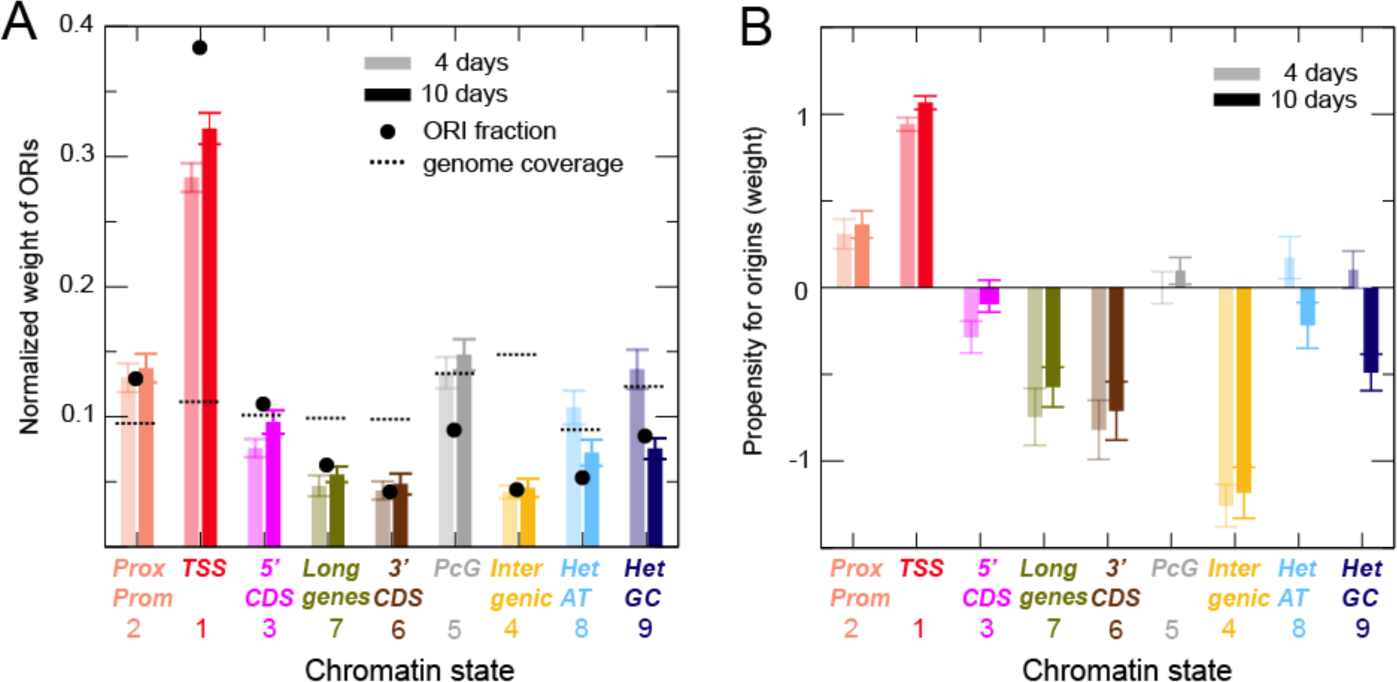
Association of ORIs with chromatin states. (A) Normalized weight of ORIs belonging to the 9 chromatin states in the combined experiments of 4 (transparent colours) and 10 day-old (solid colours) seedlings. The results of the three independent experiments are shown in the Supplemental Fig. S5A. Black circles indicate the fractions of ORIs in each chromatin state and broken lines indicate the genome coverage of each chromatin states (both using the same scale of the Y-axis). (B) Same as Fig. 4A, showing the propensity (instead of the cumulative weight) for ORIs in the 9 chromatin states is depicted. This reveals ORI types according to the associated chromatin states that have larger NSS than expected by chance based on the fraction of genome that they represent. The results of the three independent experiments are shown in Supplemental Fig. S5B.

Since not all states have the same frequency in the genome we transformed the normalized weight into propensities by dividing them by the fraction of the genome that belongs to the same state, so that positive values identify states with a NSS weight larger than expected based on its genome coverage. We confirmed that ORIs close to TSS (state 1) show the highest values, followed by ORIs in proximal promoters (state 2; Fig. 5B and Supplemental Fig. S4B). Among the rest of ORIs, those in distal regulatory intergenic regions (state 4), long genes (states 7) and 3’ end of genes (state 6) showed a negative propensity (Fig. 5B and Supplemental Fig. S4B). Interestingly, ORIs in heterochromatin (states 8 and 9) showed a negative propensity in 10 day-old but not in 4 day-old seedlings.

### Factors associated with ORI specification in different chromatin landscapes

The presence of ORIs across all the different chromatin states clearly demonstrates that ORI activity is not associated with a single chromatin signature but with different signatures depending on ORI localization. To define ORI features in a quantitative manner we calculated five traits over a region of 300 bp around the ORI midpoint for each state and experimental dataset, such as nascent strand score (NSS), CDC6, GC content, GC skew and the number of GGN trinucleotides. All of these traits correlate positively with the NSS, which suggests that they contribute positively to the strength of the ORIs. We then computed weighted averages of these traits (using the NSS as weight) over the ORI of a given state, and transformed them into Z scores, so that a positive value indicates that the trait in that state is larger than the average score over the whole genome.

We found that the GGN score profile across states is very similar to that of CDC6 and, to a lower extent, to those of GC and GC skew. Moreover, these profiles were consistently similar in all experimental situations tested (Fig. 6A and Supplemental Fig. S5), showing that these traits are rather consistent with each other. This is relevant because it might be argued that *λ*-exo has a lowered activity on G-rich secondary structures (Foulk et al. 2015). Importantly, as discussed below, both the CDC6 binding data, obtained in ChIP experiments, and ORIs mapped in Arabidopsis cultured cells, also locally enriched in GC, were obtained using independent procedures that do not rely on the use of *λ*-exo (Costas et al. 2011). We can see that ORIs in different chromatin states have different average NSS values (Fig. 6A). Thus, the average NSS of ORIs in active chromatin states 1 and 3 (TSS and 3’end of genes) is significantly lower than for ORIs in repressed heterochromatin (states 8 and 9) and Polycomb regions (state 5; the two-tailed t-test p<0.0001 in all cases except for state 3-state 9 (p<0.0046 and p<0.0002 in 10 and 4 day-old seedlings, respectively). ORIs in Polycomb chromatin were consistently high in all other traits, which can explain their high NSS. In contrast, ORIs in heterochromatin are not particularly high in the other traits. The GC content is lowest for ORIs in the distal regulatory intergenic regions (state 4), but it is still positive despite this state is AT-rich. Nevertheless, the GGN score of ORIs in state 4 is comparable to that of other states. The GGN score is the quantity that correlates most strongly with the NSS value, together with the CDC6 score (correlation coefficient 0.50 and 0.61 for the NSS of 4 and 10 day-old samples, respectively) (Fig. 6B and Supplemental Table 3). Consistent with this, the number of GGN motifs within ±150 nt around the ORI midpoint is, with some exceptions, an indication of ORI strength based on its NSS value. Furthermore, GGN trinucleotides in ORIs tend to occur in clusters larger than randomly in the average genome (Fig. 6C and Supplemental Fig. S5).

**Figure 6.**
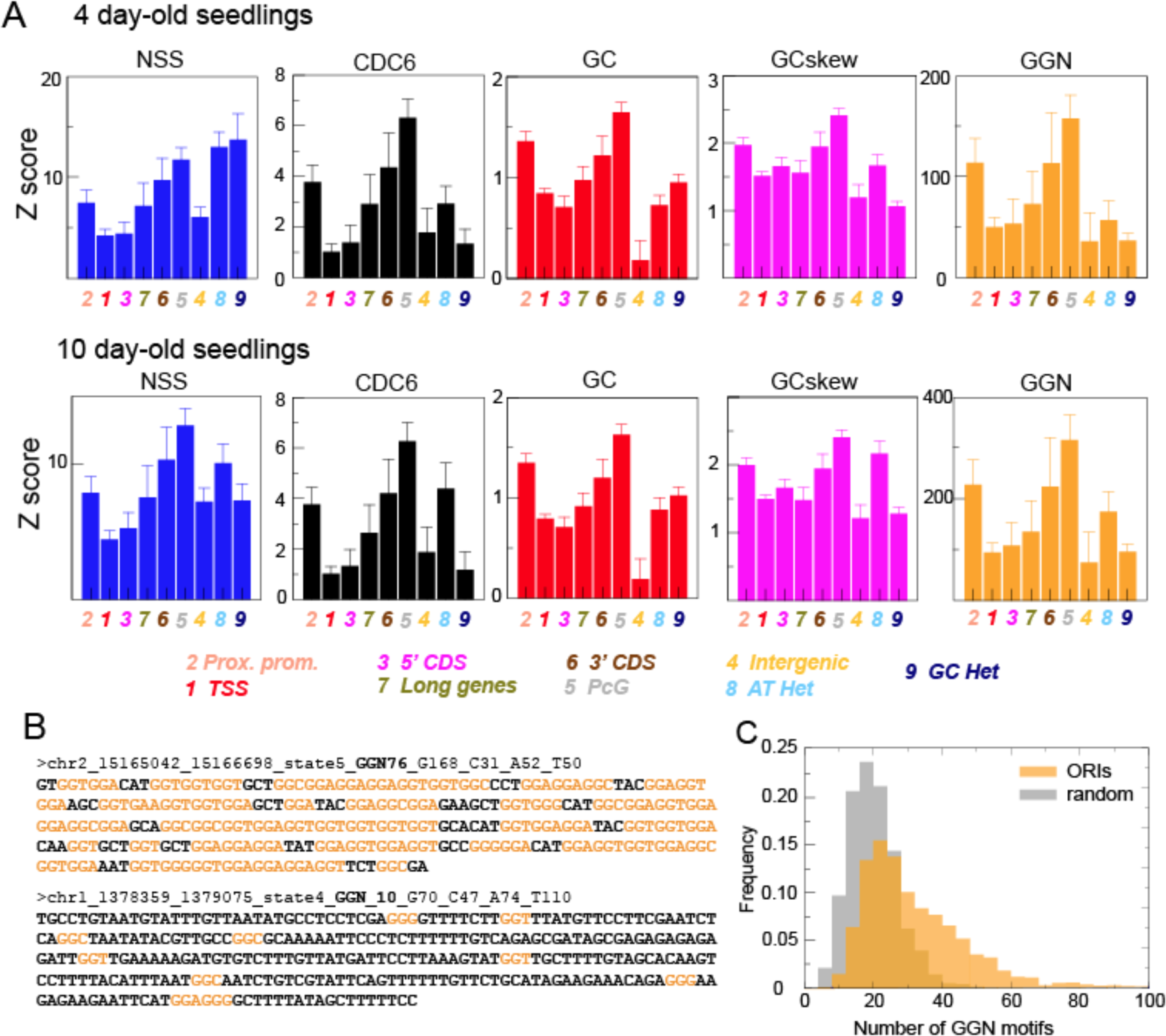
Relevance of several genomic variables for ORI specification. (A) The weighted averages of the Z scores for several variables (NSS, CDC6, GC content, GC skew and GGN trinucleotide) are shown for each chromatin state, normalized with the average property across the entire genome and weighted with the combined NSS. The results of the three independent experiments are shown in the Supplemental Fig. S6. NSS values were statistically significant when comparing states 1 and 3 with states 5, 8 and 9 (two-tailed t-test p<0.0001 in all cases except for state 3-state 9 (p<0.0046 and p<0.0002 in 10 and 4 day-old seedlings, respectively). (B) Examples of the DNA sequences ±150 nt around the ORI midpoint highlighting the GGN motifs (orange). In these two ORIs, 76 and 10 GGN motifs were present. The complete list of ORI sequences is provided in Supplemental Table 3. (C) Distribution of ORIs (n=2374) with different number of GGN motifs (orange; Mean=33.8, s.d.=17.9) compared with the same number of randomly chosen genomic regions (grey; Mean=20.6, s.d.=7.4). Two-tailed t-test, p>0.0001.

Together our results strongly support the following conclusions regarding the definition of different classes of ORIs depending on the chromatin states typical of their neighborhood.

1. ORIs located in genic regions (states 1, 3, 6 and 7), associated with active transcription and more open chromatin, possess low or intermediate NSS values, despite their large cumulative weight, suggesting a more variable usage of ORI sites. This is in part explained by the relatively lower values of CDC6 and GGN, and in part suggests interference between replication and transcription.
2. The contrary holds for ORIs in heterochromatin, with overall low accessibility for replication proteins, which tend to have a high NSS, despite their low cumulative weight. This suggested to us that once a region is specified as a potential ORI in a disfavored chromatin landscape, it is used more frequently in all cells of the population. This also applies to other compact and poorly transcribed regions such as Polycomb chromatin. It can be also considered that regions with ORIs in heterochromatin are more consistent in the different cell types leading to a stronger signal. On the other hand, active chromatin regions likely varies across cell types and ORIs associated with them are may be more variable, thus reducing the signal.

Differences in the average NSS of ORIs of different states may either stem from a global effect that affects all ORIs in the same way, or indicate strong variability of the NSS. To investigate how the chromatin states influence the variability of ORIs, we measured the correlation coefficients of the NSS over the sets of ORIs that belong to a given state (Supplemental Fig. S6). The correlation coefficients are close to one in most states, except state 1 (associated with the TSS), state 8 (AT-rich heterochromatin) and state 4 (intergenic, Polycomb repressed and AT-rich), which are most variable.

### Interplay between DNA replication origins and transcriptional programs

The relationship between ORI activity and transcriptional programs during development has been demonstrated (Nordman et al. 2011; Lubelsky et al. 2014; Comoglio et al. 2015; Muller and Nieduszynski 2017; Siefert et al. 2017). We observed that this is highly dependent on the ORI type according to the chromatin state where it is located, as visualized by plotting the quantity of transcripts in the different chromatin states identified by RNA-seq of 4 and 10 day-old seedlings. To make data comparable we transformed them into Z-scores with respect to the whole genome, so that positive values revealed more transcription than the genomic average.

The first observation is that ORI locations are more transcribed than genomic regions of the same chromatin state at both developmental stages and in all states, except GC-rich heterochromatin (state 9), supporting the strong relationship between DNA replication and transcription. Next, we compared the transcription scores of the two developmental stages. Interestingly, repressed or less transcribed regions (states 5, 4, 8 and 9) showed more transcripts through ORI sites in 4 day-old than in 10 day-old seedlings (Fig. 7 and Supplemental Fig. S7). This is particularly striking for ORIs in Polycomb chromatin (state 5) and to a lesser extent in the AT-rich heterochromatin (state 8) where a strong enhancement of transcription in 4 day-old seedlings was observed. The opposite happens for the active regions of long genes (state 7), which are more transcribed in 10 day-old seedlings. In other words, repressed regions in 4 day-old seedlings are more actively transcribed in 10 day-old, suggesting that there are differences in the accessibility of the different chromatin states, which may affect ORI activity in a developmentally regulated manner.

**Figure 7.**
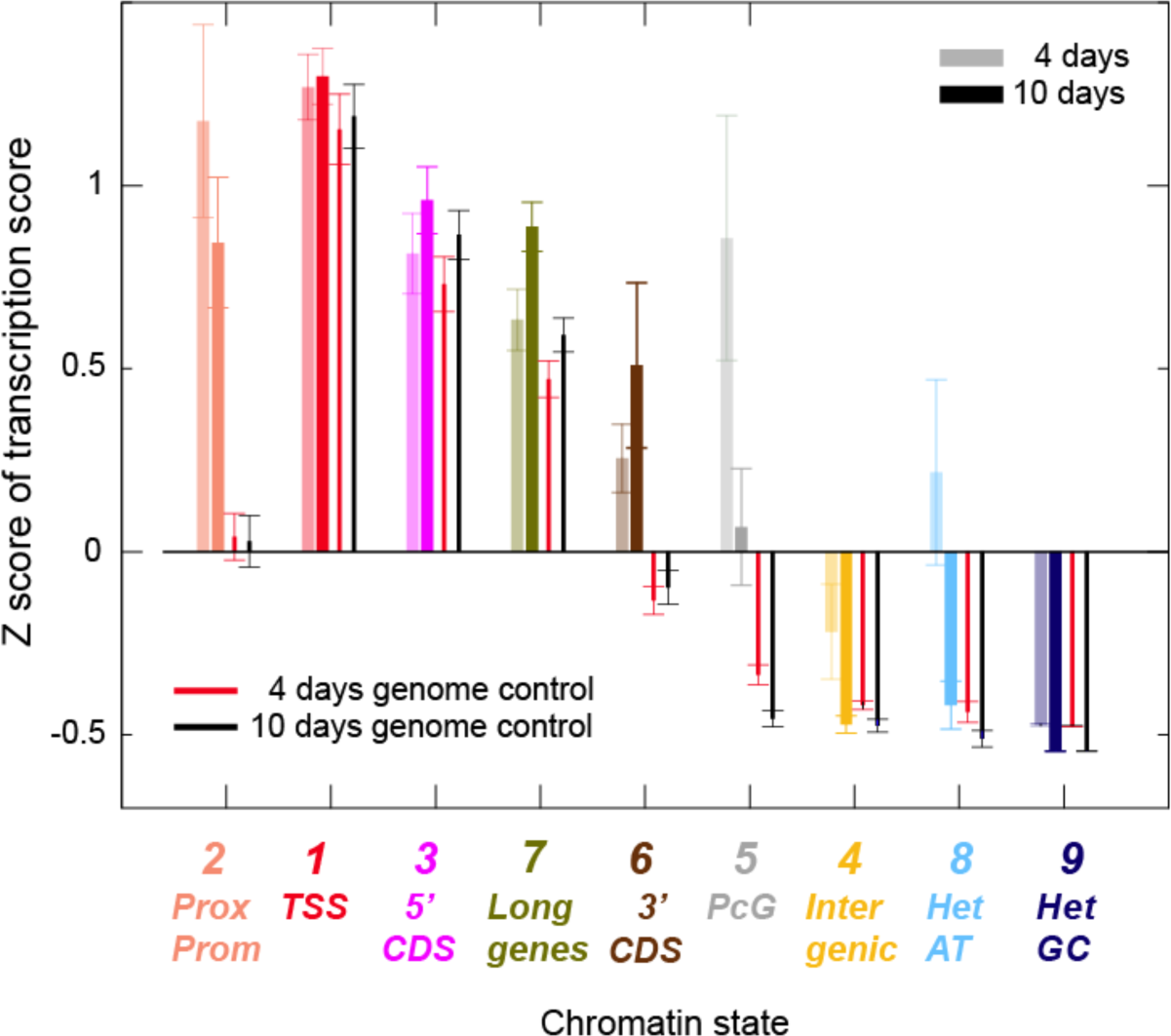
ORIs and transcriptional activity. The average Z score of the transcription score with respect to the average transcription score of the entire genome is shown for ORIs and genomic locations belonging to all chromatin states.

### ORI specification and usage during vegetative development

The systematic differences in transcriptional activity and chromatin organization observed between early (4 day-old) and late (10 day-old) vegetative stages support the idea that chromatin organization, ORI specification and the transcriptional program change during vegetative development. To further investigate these differences, we identified ORIs that have a higher NSS value in 4 day-old seedlings than in 10 day-old seedlings and vice versa, using the combined NSS of the two stages. We first analyzed the variation of the frequency of ORIs in different chromatin states as a function of the threshold used to define the preferred ORIs in each developmental stage (see Methods). To simplify the analysis we grouped ORIs in three classes: genic chromatin (states 2, 1, 3, 7 and 6), Polycomb chromatin (states 5 and 4) and heterochromatin (states 8 and 9). We found that ORIs preferentially used in 4 day-old seedlings have a strong preference for being located in heterochromatin with a very stringent threshold, whereas ORIs preferentially used in 10 day-old seedlings are located in genic states for all thresholds (Fig. 8A). Using the same threshold for both developmental times but strong enough to evidence state preferences, led us to generate a list of 94 ORIs preferentially used in 4 day-old seedlings and 91 ORIs preferentially used in 10 day-old seedlings (Supplemental Table 4).

**Figure 8.**
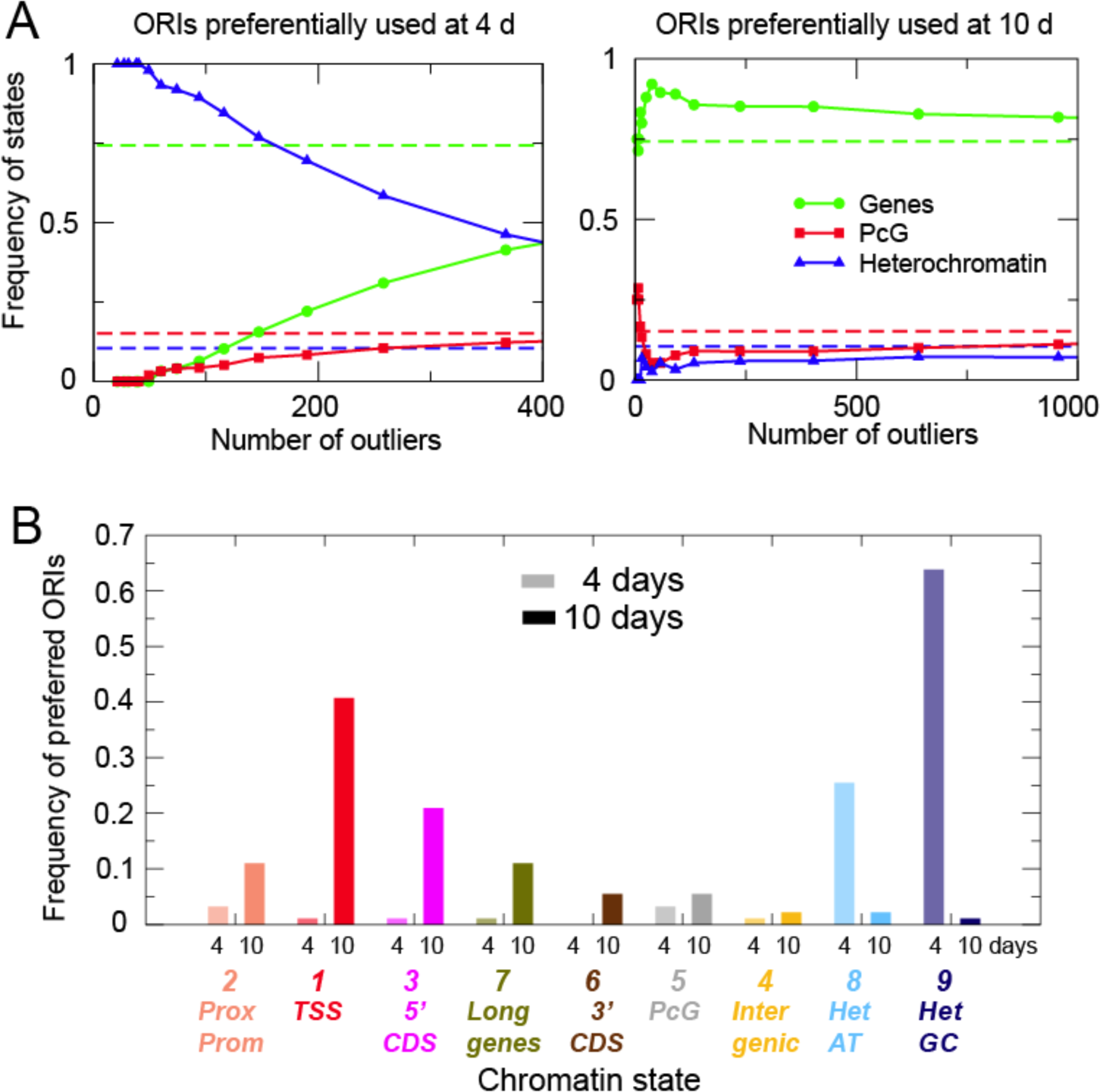
Properties of developmentally regulated ORIs. (A) Frequency of ORIs preferentially stronger in 4 day-old seedlings (left panel) or in 10 day-old seedlings (right panel), as a function of the number of ORIs obtained by varying the threshold. To simplify the analysis, ORIs have been grouped into those associated with genes (states 2, 1, 3, 7 and 6), with Polycomb chromatin (states 4 and 5) and with heterochromatin (states 8 and 9). Dotted lines represent the average values for the entire genome. (B) Frequency of ORIs preferentially used at two developmental stages defined as in panel A colocalizing with different chromatin states in 4 day-old (transparent colours) and 10 day-old seedlings (solid colours).

We analyzed several genomic features of these ORIs activated at different developmental stages. ORIs preferentially active in 10 day-old seedlings possess lower NSS, CDC6, GC and GGN scores than the average over all ORIs, indicating that they are weak (Supplemental Fig. S8). In contrast, ORIs preferentially used in 4 day-old seedlings have NSS, CDC6, GC and GGN scores comparable to or only slightly lower than the average ORIs. The most distinctive feature between these specific ORI sets is the RNA score that measures the amount of RNA reads across the regions where ORIs are located. The RNA score was comparable to the average over all ORIs for ORIs preferentially expressed in 10-day old seedlings, whereas genomic regions containing ORIs stronger in 4 day-old seedlings were significantly less transcribed.

The ORIs preferentially activated in a developmental stage-specific manner were not distributed randomly among the chromatin states. ORIs preferentially activated in 4 day-old plants occur much more frequently in the two types of heterochromatin (Fig.8B; Supplemental Table 4). Furthermore, these ORIs in TEs are more frequently located in pericentromeric heterochromatin of 4 day-old seedlings and, among them, more skewed towards Gypsy elements (81.8%), in line with data obtained in Arabidopsis cultured cells (Vergara et al. 2017). In contrast, ORIs preferentially activated in 10 day-old plants colocalize more frequently with genic regions, typically in 5’-end of genes, TSS and proximal promoters (Fig. 8B). Since ORI activation is highly related to chromatin accessibility, our data strongly suggest that ORI usage changes significantly in a locus-specific manner during postembryonic development, most likely associated with changes in chromatin organization.

## Discussion

### Genomic features of Arabidopsis DNA replication origins

In this study, we have generated a whole-body originome map with the location and properties of ORIs in *A. thaliana* plants at two different developmental stages of vegetative growth. ORI activity was assessed quantitatively from the sequencing data by determining a nascent strand score (NSS) that measures the propensity of a certain genomic location to behave as an ORI.

We have identified 2374 potential ORIs in the Arabidopsis genome that can function during vegetative growth. Our results indicate that Arabidopsis ORIs are organized in discrete sites rather than in large initiation zones, in agreement with ORI mapping in cells with similar genome size (Comoglio et al. 2015; Lombrana et al. 2016; Pourkarimi et al. 2016; Rodriguez-Martinez et al. 2017). A comparison of ORI locations in cultured cells and seedlings revealed a coincidence of ~14-25%, depending on the threshold tolerance. This amount of ORIs common to both sources reveals the existence of technical and biological variables in ORI usage, e.g. the presence of many different cell types in the seedling compared to the cell culture, that need to be identified in the future using the tools established in this work.

Most Arabidopsis ORIs in seedlings (~78%) associate with genic elements, in particular the 5’ end of genes, reinforcing the strong preference of ORIs for genic regions. Overall this is similar to the situation in metazoan cultured cells and embryos (Sequeira-Mendes et al. 2009; Macalpine et al. 2010; Cayrou et al. 2015; Comoglio et al. 2015; Rodriguez-Martinez et al. 2017). Similar to the situation in cultured cells (Costas et al. 2011), Arabidopsis ORIs colocalize with transposable elements (TEs) less frequently than expected at random. However, as previously reported in cultured cells (Vergara et al. 2017), we found that ORIs in pericentromeric regions, which contain a lower gene density, increase their tendency to colocalize with TEs and in particular with retrotransposons of the Gypsy and LINE families. These results reinforce the conclusion that Arabidopsis ORIs have a high preference of being associated with genes in euchromatin and with both genes and transposons in pericentromeric heterochromatin. Moreover, we show that in seedlings the frequency of ORIs colocalizing with DNA transposons, mostly of the MuDR family, is much higher that in cultured cells, perhaps due to a different chromatin landscape in cultured cells and seedlings.

#### GGN clusters are a strong determinant of ORI strength

The number of GGN trinucleotides within ±150 nt around the ORI midpoint is the feature that correlates most strongly with the NSS value, although it explains <30% of the variance (r^2^=0.28). From a structural point of view, four consecutive GGN motifs may form G-rich secondary structures, such as G4 with two tetrads (Sen and Gilbert 1988; Chen and Yang 2012). Although some computer programs require at least three tetrads to identify G4 (Todd et al. 2005), examples with two tetrads have been experimentally characterized, such as the thrombin binding aptamer d(G2T2G2TGTG2T2G2) (Macaya et al. 1993) and the *Bombyx mori* telomeric sequence d(AG2T2AG2T2AG2T2AG2) (Sacca et al. 2005). Moreover, the stability of G4 is largest for loops of length 1, such as those in consecutive GGN motifs, and it has been observed that GjNGj sequence motifs form a robust parallel stranded structure motif with 1 nt loop (Chen and Yang 2012). Stabilizing interactions between distinct G4 structures have been observed (Palumbo et al. 2009), suggesting that there could be synergy between the large number of GGN motifs observed in Arabidopsis ORIs.

Only 35 (1.4%) ORIs contain <4 GGN motifs, while the maximum frequency is between 8 and 16 GGN, finding up to 77 GGN out of the theoretical maximum of 100 in the 300 nt windows. Such nucleotide distribution produces one strand very enriched in Gs, favoring the possibility to form G4 (Cayrou et al. 2015). Our finding that GGN motifs are enriched in ORIs is consistent with reports that the presence of G4 influences ORI activity in animal cells (Besnard et al. 2012; Cayrou et al. 2012; Valton et al. 2014; Cayrou et al. 2015). Moreover, the local GC content is much higher for ORIs than for random genomic locations, which seems to be a common feature of ORIs in all animal and plant cells mapped so far (Macalpine et al. 2010; Cayrou et al. 2011; Costas et al. 2011; Besnard et al. 2012; Cayrou et al. 2015). The enrichment of ORIs in GGN motifs might be related to a reduced efficiency of X-exo to digest G-mediated secondary structures. However, we have carried out *λ*-exo treatments under optimal enzyme/substrate conditions. Moreover, it must be kept in mind that (1) we observed a strong correlation between occurrence of GGN motifs at ORIs and the CDC6 binding score, and (2) Arabidopsis ORIs were found to be locally enriched in Gs in cultured cells using procedures that do not rely on *λ*-exo treated samples (Costas et al., 2011).

The enrichment in GGN stems from two genomic properties of Arabidopsis ORIs, the high GC content and the GC skew, which concur to produce one G-rich strand. As mentioned above, the GC skew associated with replication has been prevalently associated with the mutational asymmetry between the leading and lagging strand (Lobry 1996). However, we found that the asymmetry between G and C starts approximately 300 nt before the ORI, identified as the local maximum of the NSS. Thus, the replication asymmetry cannot be the sole cause of the GC skew at Arabidopsis ORIs, which may be likely generated by positive selection of GGN clusters. Moreover, many GGN clusters are formed by tandems of quasi-repeats, so that a likely mutational mechanism is through trinucleotides insertions produced by the slippage of the polymerase. Importantly, the three hypothesized mechanisms, replication asymmetry, insertion and selection, are expected to cooperate for the formation and maintenance of GGN clusters.

#### Other structural features of Arabidopsis DNA replication origins

Besides GGN clusters we have identified other genomic features associated with the ORI activity. A first important feature is chromatin accessibility because more accessible states, which are transcriptionally active, are more prone to be ORI locations. The second driving factor is the propensity to bind the pre-initiation protein CDC6. Interestingly, the affinity for CDC6 is on average significantly larger in the repressed than in the active chromatin states, probably in order to compensate their reduced accessibility. Finally, despite both transcription and replication activities are affected by chromatin accessibility, we observed an overall negative correlation between the average ORI strength and transcription of the chromatin state. In particular, ORIs colocalizing with TSS are overall among the weakest ones. There are reports of a preferential location of ORIs in actively transcribed genes (Aladjem 2004; MacAlpine et al. 2004; Saha et al. 2004; Goren et al. 2008). However, in these cases the ORIs are not in close proximity to the TSS. We hypothesize that the negative correlation may originate from possible interference between replication and transcription (Aguilera and Garcia-Muse 2013).

Our data clearly demonstrate the existence of different types of ORIs according to their features, including primarily their chromatin landscape. There are multiple signatures that can accommodate ORIs to ensure full replication of the entire genome, although ORIs show an overall preference for localizing in the TSS (state 1) and adjacent states containing relatively open chromatin properties. This points to the compatibility of ORI activity with multiple chromatin signatures (Cayrou et al. 2015). Long genes (state 7), the 3’-end of coding sequences (state 6) and the distal regulatory intergenic regions (state 4) are significantly depleted of ORIs. Surprisingly, ORIs located within PcG repressed chromatin (state 5) as well as within heterochromatic domains (states 8 and 9) tend to be stronger than average. Thus, it is conceivable that finding an appropriate local ORI landscape within repressed and compact chromatin may favor that it is used in more cells of the population, leading to a higher NSS value.

#### Developmental regulation of DNA replication origins and the chromatin landscape

We found remarkable that ORI activity undergoes systematic changes in the course of development. In particular, we observed that ORIs that are stronger in 10 day-old seedlings are particularly frequent in the TSS (state 1) and adjacent chromatin regions (states 2 and 3). In contrast, ORIs preferentially used in 4 day-old seedlings are particularly recurrent in heterochromatin (state 9). These trends are consistent with our transcriptional analysis that revealed lower repression of typically repressed chromatin states in 4 day-old seedlings compared to 10 day-old seedlings, suggesting that chromatin organization may be different at these two stages of vegetative development. The increased frequency of ORIs in TEs, in particular of the Gypsy family, in 4 day-old seedlings suggests that the role of these TEs as ORIs could be important at early developmental stages where the dynamics of cytosine methylation suggests differences in the repression level of heterochromatin (Bouyer et al. 2017). This is a major difference with recent studies in *C. elegans*, where early pregastrula embryos are depleted of ORIs in heterochromatin whereas ORIs have a preference for non-coding regions and enhancers in postgastrula embryos (Rodriguez-Martinez et al. 2017). These differences between animal and plants suggest different mechanisms of coupling ORI activity, developmental programs and heterochromatin dynamics. It is also worth noting that the spatial organization of the Arabidopsis genome reveals that typical TADs and distal enhancers as in animals are lacking, or very infrequent (Wang et al. 2015; Liu et al. 2016; Vergara and Gutierrez 2017).

Based on ORI identification in various cell types in culture, a general consensus exists that developmentally regulated ORIs are not very efficient (Besnard et al. 2012). The quantitative parameter of ORI activity (NSS), developed in our study, clearly showed that every genomic location associated with an ORI possesses a certain firing efficiency. In agreement with studies in animal cells in culture (Besnard et al. 2012; Comoglio et al. 2015), it seems that modulation of ORI activity rather than the selection of different genomic locations determines ORI usage at different developmental stages and, most likely, in different cell types.

Polycomb complexes are involved in regulating gene expression associated with developmental phase transitions in Arabidopsis (Kuwabara and Gruissem 2014). We found ORIs associated with Polycomb-regulated genes (state 5), although they are under-represented in these chromatin regions. Somehow surprisingly, they behave as strong ORIs, suggesting that once a genomic site is chosen as an ORI within a Polycomb-repressed chromatin domain, the position of ORIs is restricted and used in many cells. This is in agreement with the hypothesis developed for mammalian cells in culture, where Polycomb factors are strongly associated with efficiently used ORIs (Cayrou et al. 2011; Picard et al. 2014; Cayrou et al. 2015). Developmentally regulated genes in animal pluripotent stem cells share H3K4me3 and H3K27me3 marks, typical of bivalent chromatin (Bernstein et al. 2006). An attractive possibility is that some of the H3K27me3 regions, colocalizing with ORIs, may actually constitute regions of bivalent chromatin, although this has not been experimentally demonstrated. Using sequential re-ChIP experiments (Sequeira-Mendes et al. 2014) we demonstrated the presence of this bivalent chromatin type in somatic cells of Arabidopsis seedlings, restricted to proximal and distal regulatory regions (states 2 and 4, respectively). We also found that H3K27me3 (and H3K4me3) is enriched in ORIs located in proximal promoters (state 2) but not in distal regulatory regions (state 4). Thus, it is conceivable that the subset of ORIs potentially associated with bivalent chromatin may have a fast response to developmental cues, as it occurs to genes where they colocalize.

Our genome-wide results have defined the main DNA and chromatin properties associated with different ORI classes in a living organism and at two growth stages during postembryonic vegetative growth. These properties allowed us to show that ORI activity is compatible with a variety of signatures and demonstrate the existence of various classes of ORIs defined by their strength, DNA features and chromatin landscape. The feasibility to study ORI activity in a developmentally and genetically tractable organism opens new avenues to determine how ORI activity is regulated in response to developmental cues, in association with transcriptional programs, in response to environmental challenges and in a variety of mutant backgrounds.

### Methods

#### Plant growth

*Arabidopsis thaliana* seeds (Col-0 ecotype) were stratified for 48h and grown in Murashige and Skoog (MS) medium supplemented with 1% (w/v) sucrose and 1% (w/v) agar in a 16h:8h light/dark regime at 22°C, for either 4 or 10 days.

#### Purification of short nascent strands (SNS)

Total genomic DNA and SNS preparations were obtained under RNase-free conditions, by an optimization of the protocol described (Sequeira-Mendes et al. 2009), which has been enhanced to obtain sufficiently clean SNS preparations. Nuclei isolation from 4 or 10 days post-sowing (dps) Arabidopsis seedlings was performed prior to genomic DNA extraction as described (Chodavarapu et al. 2010), in order to minimize genomic DNA contamination with cytosolic polyphenols and other secondary metabolites. Twelve grams of whole seedlings were collected, frozen and ground in liquid nitrogen in the presence of 10% PVPP (Sigma). The ground material was resuspended in 10 ml per gram of Honda Buffer Modified for 30 min in a rotary shaker at 4 °C (HBM; 2% (p/v) PVP10 (Sigma), 25 mM Tris-HCl, pH 7.6, 440 mM sucrose (Merck), 10 mM magnesium chloride, 0.1% Triton X-100, 10 mM β-mercaptoethanol). To better release the nuclei, the resuspended material was processed in a dounce homogenizer twice with a loose and a tight pestles and filtered through a double miracloth mesh into corex tubes. The nuclei were centrifuged 10 min at 3000xg and 4 °C. The supernatant was discarded and nuclear pellet was resuspended in 5 ml per gram of Nuclei Isolation Buffer (NIB; 2% (p/v) PVP10 (Sigma), 20 mM Tris-HCl, pH 7.6, 250 mM sucrose (Merck), 5 mM magnesium chloride, 5 mM potassium chloride, 0.1% Triton X-100, 10 mM β-mercaptoethanol). The sample was loaded onto a 15/50% gradient of percoll in NIB and centrifuged 20 min at 500x*g* and 4 °C with slow brake. The green upper layer was discarded and the same volume was replaced with NIB. Nuclei were centrifuged 5 min at 1100x*g* and 4 °C, washed twice with 10 ml of NIB and 4 °C, and resuspended in 20 ml of lysis buffer per 12 grams of starting material (0.5% (p/v) PVP10, 50 mM Tris-HCl, pH 8.0, 10 mM EDTA pH 8.0, 1% SDS, 10 mM β-mercaptoethanol) by agitation 15 min at 4 °C. To digest the proteins, 100 μg/ml proteinase K was added and incubated overnight at 37 °C with mild rotation. Total DNA was extracted twice, first using phenol, pH 8.0, then with phenol:chloroform:IAA and the aqueous phase containing genomic DNA was collected into polyallomer tubes (Beckman). DNA was precipitated by adding 1.5M sodium chloride and 2 volumes of absolute ethanol, incubated 1h at −80 °C and pelleted by centrifugation for 45 min at 52,000x*g* at 4 °C using an AH-627 rotor (Sorvall). DNA was washed twice with 70% ethanol, centrifuged 20 min at 52,000xg and room temperature in an AH-627 rotor (Sorvall), air dried and resuspended in 1 ml of TE (10 mM Tris-HCl, pH 8.0, 1 mM EDTA) containing 160 U of RNase OUT (Invitrogen). DNA was incubated at 4 °C overnight without pippeting or vortexing.

Purified DNA was denatured by heating 10 min at 100 °C and size-fractionated in a seven-step neutral sucrose gradient (5-20% sucrose in TEN buffer (10 mM Tris-HCl, pH 8.0, 1 mM EDTA and 100 mM sodium chloride), by centrifugation at 102,000xg in a SW-40Ti Beckman rotor for 20 h at 20 °C (Gomez and Antequera 2008). Fractions (1 ml) were collected from the top and the DNA was ethanol-precipitated. An aliquot of each fraction was analyzed in a 1% alkaline agarose gel (50 mM sodium hydroxyde, 1 mM EDTA) to monitor size fractionation. Normally, fractions 3 (~100-600 nt), 4 (~300-800 nt) and 5+6+7 (~500-3000 nt) were processed further by treating with 0.67 U/μl of polynucleotide kinase (PNK, Fermentas) to phosphorylate 5’-hydroxyl ends in the presence of 1.34 mM dATP for 30 min at 37 °C. After PNK inactivation, phosphorylated DNA was extracted, precipitated and resuspended in water. SNS were distinguished from randomly broken DNA molecules based on the presence of 4-6 nt-long RNA primers at their 5’-ends, which made them resistant to λ-exonuclease treatment (Gerbi and Bielinsky 1997; Costas et al. 2011; Cayrou et al. 2015; Comoglio et al. 2015). The λ-exonuclease digestion was carried out with 5 U/μl of enzyme (Thermo Scientific) following the manufacture’s instructions at 37 °C overnight. The efficiency of the digestion was monitored by adding 40 ng of phosphorylated linearized plasmid to an aliquot of each reaction tube. DNA from each λ exonuclease-treated fraction was extracted, precipitated and resuspended in TE. The phosphorylation and λ-exonuclease treatments were repeated at least twice. RNA was digested with 0.05 μg/ml RNase A (Roche) and 0.16 U/μl RNase I (Thermo Scientific) for 30 min at 37 °C. RNases were digested with 100 μg/ml proteinase K and DNA was extracted, precipitated and resuspended in miliQ water. The ssDNA of purified SNS was converted into dsDNA: first, SNS together with 2 pmol random hexamer primers (Roche) were denatured 5 min at 100 °C, then a slow annealing was achieved by cooling down the samples from 80 °C to room temperature; second, the dsDNA was synthesized by using 0.17 U/μl of Klenow fragment for 1h at 37 °C; third, the fragments were ligated with 2 U/μl of Taq DNA ligase (New England Biolabs) for 45 min at 45 °C; finally, dsDNA was extracted, precipitated, resuspended in miliQ water and quantified before proceeding to the library preparation. The same method of dsDNA conversion was applied to sheared and denatured genomic DNA to be used as sequencing control.

#### RNA purification

Total RNA from 4 pds seedlings was isolated using Trizol (Invitrogen) according to the manufacturer’s instructions. Total RNA was treated with DNase I (Roche) before proceeding with library preparation.

#### Next-generation sequencing

DNA libraries of both SNS DNA (sucrose gradient fractions 3, 4 and 5+6+7 combined) and gDNAs were first sheared by an S2 focused-ultrasonicator (Covaris) for 2 minutes (Intensity 5, Duty Cycle 10%, Cycles per Burst 200), and then used as inputs to generate sequencing libraries by Ovation Ultralow V1 library prep kits (NuGen). The libraries were subjected to deep sequencing on HiSeq 2000 per manufacturer instructions (Illumina). In two out of the three experiments we used different amplification protocols for library generation with the purpose of estimating possible bias introduced by this crucial but unavoidable step. RNA-Seq libraries were made by TruSeq Stranded mRNA library prep kit and NeoPrep (Illumina), and subjected to deep sequencing on HiSeq 2000 per manufacturer instructions (Illumina). Single-end sequenced reads (51 nt) were aligned to the reference Arabidopsis genome (TAIR10), using the Bowtie alignment tool (Langmead et al. 2009), allowing up to one mismatch and discarding multihit reads. PCR duplicate reads were removed using an in-house script.

#### Peak-calling

For each sample and each fraction, we call ORIs with our own peak calling algorithm ZPeaks (Bastolla et al., in preparation) that (i) provides a well defined, genome-wide profile of Nascent Strand Score (NSS), instrumental for weighting candidate ORIs and generic genomic locations, and (ii) localizes an ORI at the local maximum of the NSS over the ORI box called, needed for centering the metaplots. We tested ZPeaks by visual inspection of the overlap between experiment and control reads and candidate ORIs as well as by the statistical analysis of the ORIs properties. Furthermore, our procedure was robust with respect to false positive ORIs because (i) it requires that each ORI is detected in several independent experiments and (ii) it weights each ORI with its NSS, so that spurious ORIs have low NSS and contribute little to the average properties.

Thus, ZPeaks computes optimally smoothed profiles of the reads of the experiment and the control, obtains from them a normalized smoothed profile, calls peaks when the profile is above an user-specified threshold, and sets the ORI location at the maximum of the normalized profile. More in detail, the algorithm works as follows, once the sequencing reads have been aligned to the reference Arabidopsis TAIR10 genome: (1) The wig files (normalized read counts) are input to ZPeaks and the number of reads is rescaled so that its mean number over each chromosome is the same both for the experiment *e* and the control *c*. If the control is not available, a constant profile is used. To increase the reliability of bins where the control is low, values of the control below the mean are interpolated between the current value and the mean: if *c_i_* < <*c*>, then we use *c_i_*′ =(*c_i_*+<*C*>)/2, where *i* indicates the genomic location and <*C*> is the mean value of *C* over the chromosome where *i* is located; (2) The profiles of the rescaled experiment and control are smoothed as *c*_*i*_^′^=*Σ*_*k*_ *c_i_ w_ik_* where the weights *w_ik_* are given by *w_ik_*=*exp(*-*d_ik_* /*d*_0_)/*Σ_l_ exp(-d_il_ /d_0_)*, *d_ik_* is the distance between the center of bin *i* and bin *k*, *d_0_* is a parameter that is optimized as described below. A cut-off on distance is used to accelerate the computation, whose value is optimized alongside d0; (3) From the smoothed experiment and control, the difference score *d_i_* = *e_i_*′-*c_i_*′ is constructed and it is transformed into the Z score *z_i_* = *(d_i_* −<*d*>)/*s_d_*, where <d> is the mean value of *d* over the chromosome of *i* and *s_d_* is the standard deviation; (4) For the chosen threshold *T*, the program counts the number of bins with *z_i_* > *T*, *N(T)*. The smoothing parameter *d_0_* that yields the largest *N(T)* for the chosen threshold is chosen as the optimal parameter. The rationale is that, if the profiles are smoothed too much, then the experiment and the control will tend to become equal to their mean values and *d_i_*, will tend to be zero, thus decreasing N(T), whereas if the profiles are smoothed too little the standard deviations will be large, also decreasing N(T). We can always determine numerically an optimal parameter *d_0_* for which *N(T)* is maximum, which justifies our procedure. We defined the nascent strand score (NSS) profile of the experiment *e* as NSS_*ei*_ = *z_i_*; (5) We then joined together consecutive bins with NSS_*ei*_ > T separated by less than 200 nucleotides, obtaining boxes that represent candidate origins; (6) Finally, the putative DNA replication origin is set at the bin where *z_i_*, is maximum within the box, and the limits of the box are reduced in such a way that the ORI is at the center and the new box is contained into the original one. It must kept in mind that the NSS showed a continuous distribution without any sign of saturation, perhaps suggesting that some bias introduced by the amplification step prior to sequencing may have some contribution.

One may expect that the threshold parameter T may be objectively determined by clustering all genomic bins in two clusters through some clustering algorithm such as K-means, Expectation Maximization (that assumes that the scores *z_i_* are distributed according to a Gaussian distribution) or Hidden Markov Models (that also exploits the positional order of the bins along the chromosome). We followed such strategies, but the thresholds that we obtained were low, a sizable fraction of the genome satisfied *z_i_* > *T*, and visual inspection showed that most candidate ORIs were not reliable. Thus, we had no better choice than selecting an arbitrary threshold T and determining *bona fide* ORIs by combining different experiments, as explained below.

#### Combining peaks into consensus boxes (potential ORIs)

Our strategy consisted of determining a robust set of ORIs detected in at least two independent experiments and two fractions for each experiment and weighting each candidate ORI with the NSS value of each experiment in such a way that the results are little dependent of false positives with low score.

We analyzed two developmental stages (4 and 10 day-old seedlings) and 3 experiments for each stage (exp1, exp2, exp3), obtaining six different samples. For each of them, either two (F3 and F4) or three (F3, F4, F5+6+7) consecutive fractions of the sucrose gradients for size selection of nascent strands were sequenced. Fractions were equivalent between 4 and 10 day-old seedlings, matching F3, F4 and F5+6+7. We called candidate ORIs with a tolerant threshold (z>1.8) and for each sample we selected candidate regions, or boxes, that were identified in at least two fractions of the same gradient. Boxes with size smaller than 200 bp were eliminated, and boxes closer than 200 bp were joined. In this way we obtained six datasets of high quality ORIs, which numbers were: 842 (4d_exp1), 1938 (4d_exp2), 3008 (4d_exp3), 3298 (10d_exp1), 1686 (10d_exp2), 3107 (10d_exp3).

To increase the reliability of candidate ORIs, we selected only those boxes that had been found in at least two out of six independent samples, obtaining a total of 2374 highly reliable candidate ORIs. We matched the boxes with non-vanishing overlap and if an ORI had multiple overlaps, we selected the largest overlap. The center of the combined box was computed as the weighted average of the location with maximum score present in the associated boxes, weighting more the boxes with high NSS and small size. When we matched different fractions, the fraction F5+6+7, which contains larger nascent strands, was used to confirm boxes but not to locate their center, in order to obtain better resolution. The limits of the combined box were set in such a way that all of the bins are above the threshold in all fractions.

#### Scoring ORIs in different samples

For each ORI, we obtained their score NSS_*ek*_ in the six samples, where *e* labels the experiment, *k* labels the ORI, and NSS_*ek*_ is the maximum value of the score over all bins included in the box that contains the ORI. For each sample, we used the corresponding scores as weights, and we obtained the average values and the metaplots of genomic and epigenetic marks as the weighted average over the set of ORIs. We also generated combined scores by averaging the scores of all 4 day-old and all 10 day-old seedling samples.

#### Quantitative real-time PCR (qPCR)

The qPCR analysis was performed using GoTaq Master Mix (Promega) according to the manufacturer’s instructions in an ABI Prism 7900HT apparatus (Applied Biosystems). For each region a subset of unique and specific primers, listed in Supplemental Table 2, were designed. The quantification was determined using a standard curve (five serial 4-fold dilutions of gDNA) and SNS enrichment was normalized against a region lacking ORIs (negative control).

#### Detection of developmentally regulated ORIs

Despite most ORI scores were strongly correlated, indicating that ORIs that are strong in one sample are strong also in the others, we identified a reduced number of ORIs whose strength is significantly different from one sample to the other. For this purpose, we only considered the combined scores of the two developmental stages (4 and 10 day-old seedlings), we rescaled them in such a way that the average value over the set of origins was the same for both samples, and for each ORI *k*. We computed the mean and standard deviation of the rescaled NSS_*ek*_ over the samples. We considered variable those ORIs in which the ratio between the standard deviation and the mean is larger than a threshold and studied the properties of outliers as a function of the threshold.

Analysis of ORIs in heterochromatin and TE families was carried out as described in (Vergara et al. 2017).

## Data access

The SNS-seq datasets generated in this study have been submitted to the NCBI Gene Expression Omnibus (GEO; http://www.ncbi.nlm.nih.gov/geo/) under accession number GSE109668.

## ACKNOWLEDGMENTS

We thank E. Martinez-Salas and M. Gomez for discussions and comments during this project and the critical reading of the manuscript, C. Becker and N. Schandry for useful suggestions, members of our laboratories for continuous feedback and V. Mora-Gil for technical assistance. We thank S. Jacobsen, C. Hale, S. Feng and the BSCRC BioSequencing Core for Illumina DNA Sequencing and discussions. This research was supported by grants BFU2012-34821, BFU2013-50098-EXP and BFU2015-68396-R to C.G., BIO2016-79043-P to U.B. and AGL2013-43244-R to J.M.C., as well as by institutional grants from Fundacion Ramon Areces and Banco de Santander to the CBMSO.

## Author contributions

The work was conceived by CG, JS-M and UB. JS-M, together with ZV, implemented protocols for purification and analysis of nascent strands, with the help of CC and IA in the initial steps of the work. UB developed the computational and statistical analysis, with the initial help of RM-G. JS-M, ZV, UB and RP generated and analyzed data. JM and JMC analyzed ORIs in heterochromatin. CG and UB wrote the manuscript with the input of all authors.

## Supplemental Material

Supplemental material includes eight Supplemental Figures (S1-S8) and four Supplemental Tables.

## SUPPLEMENTAL Material

### Supplemental Figures

**Figure S1.**
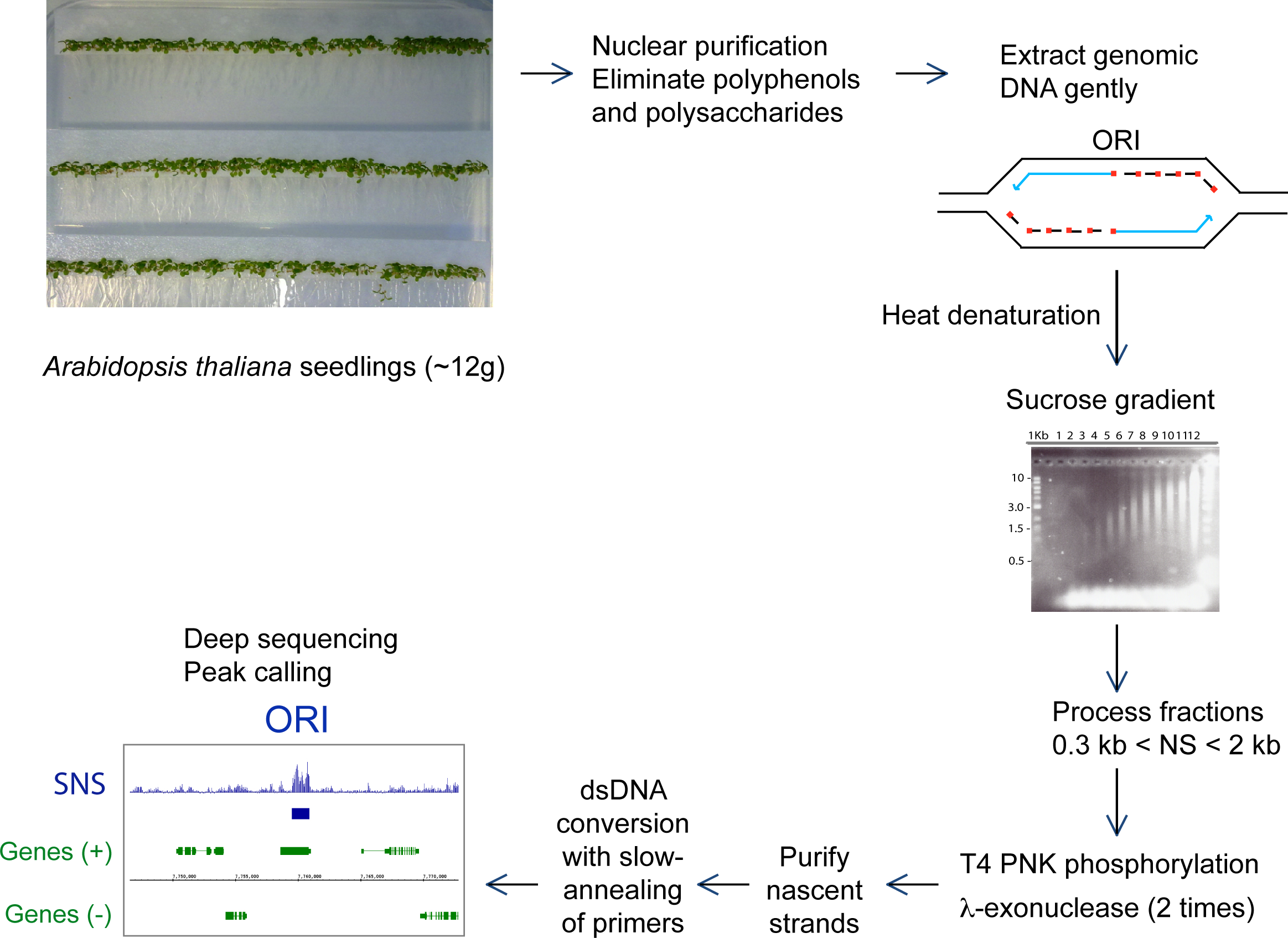
Enhanced protocol for purification of nascent strands (NS) from whole developing seedlings.

**Figure S2.**
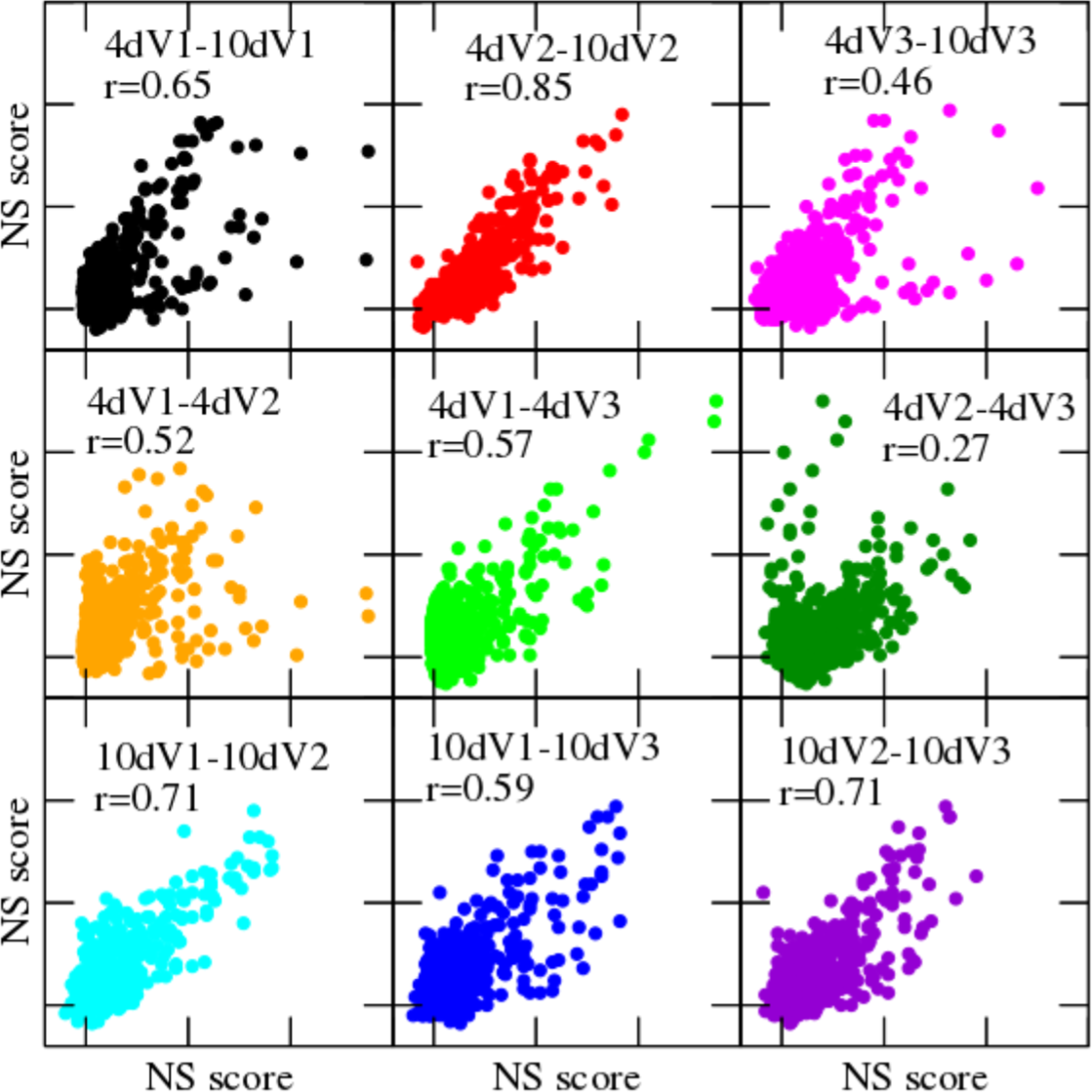
Scatter plots of the nascent strand score (NSS) measured in different experiments. Each point represents one of the 2374 ORIs identified in this study. Only pairs corresponding to the same day or the same experiment are shown. V1, V2 and V3 refer to each of the three experiments.

**Figure S3.**
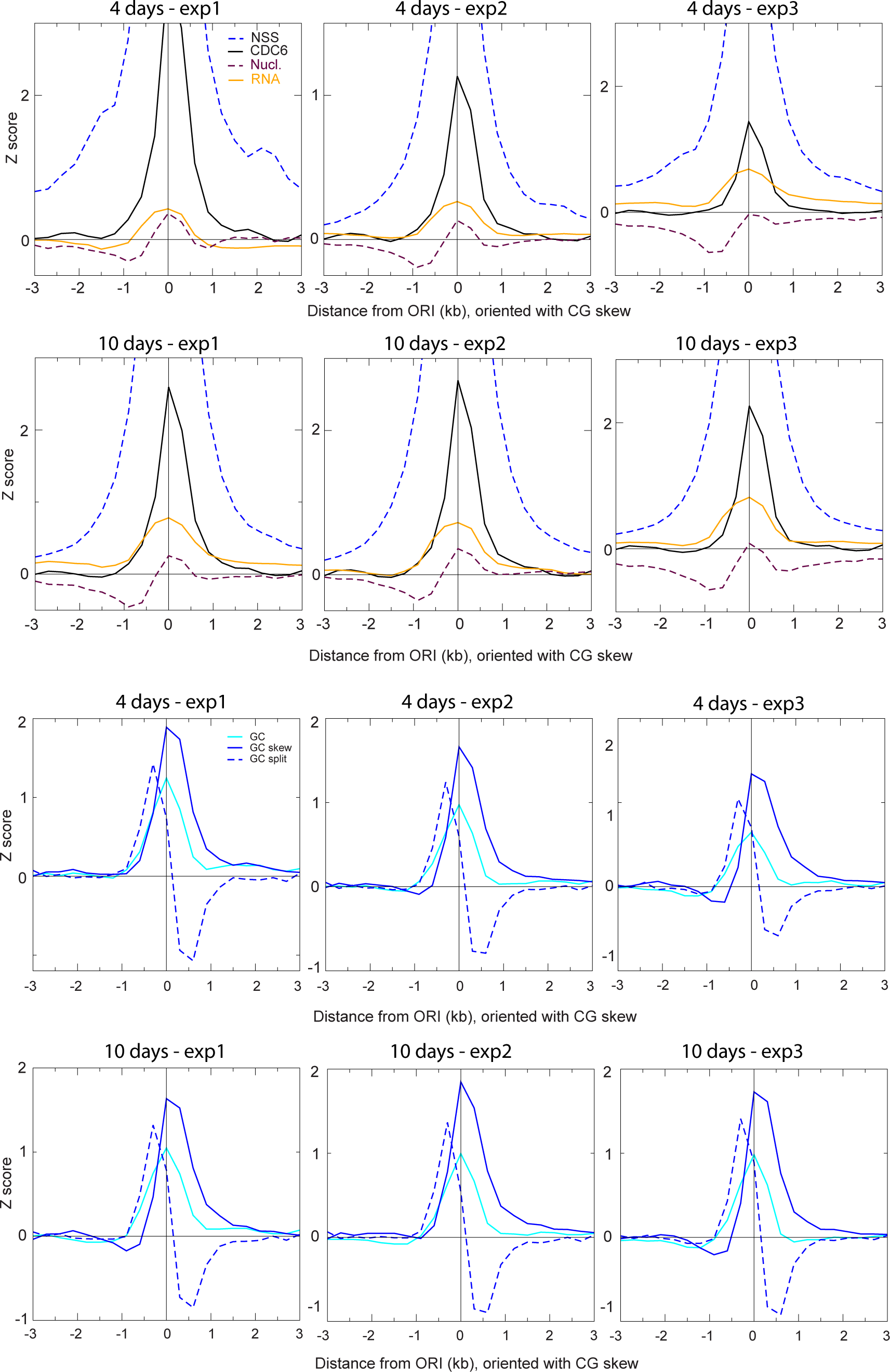
Features of the local neighborhood of ORIs in whole Arabidopsis seedlings. Metaplots of NSS, CDC6, transcript content (RNA) and nucleosome (nucl.) content (upper two panels) and of GC, GC skew and GC split (lower two panels) of each of the three independent experiments in 4 and 10 day-old seedlings, as indicated.

**Figure S4.**
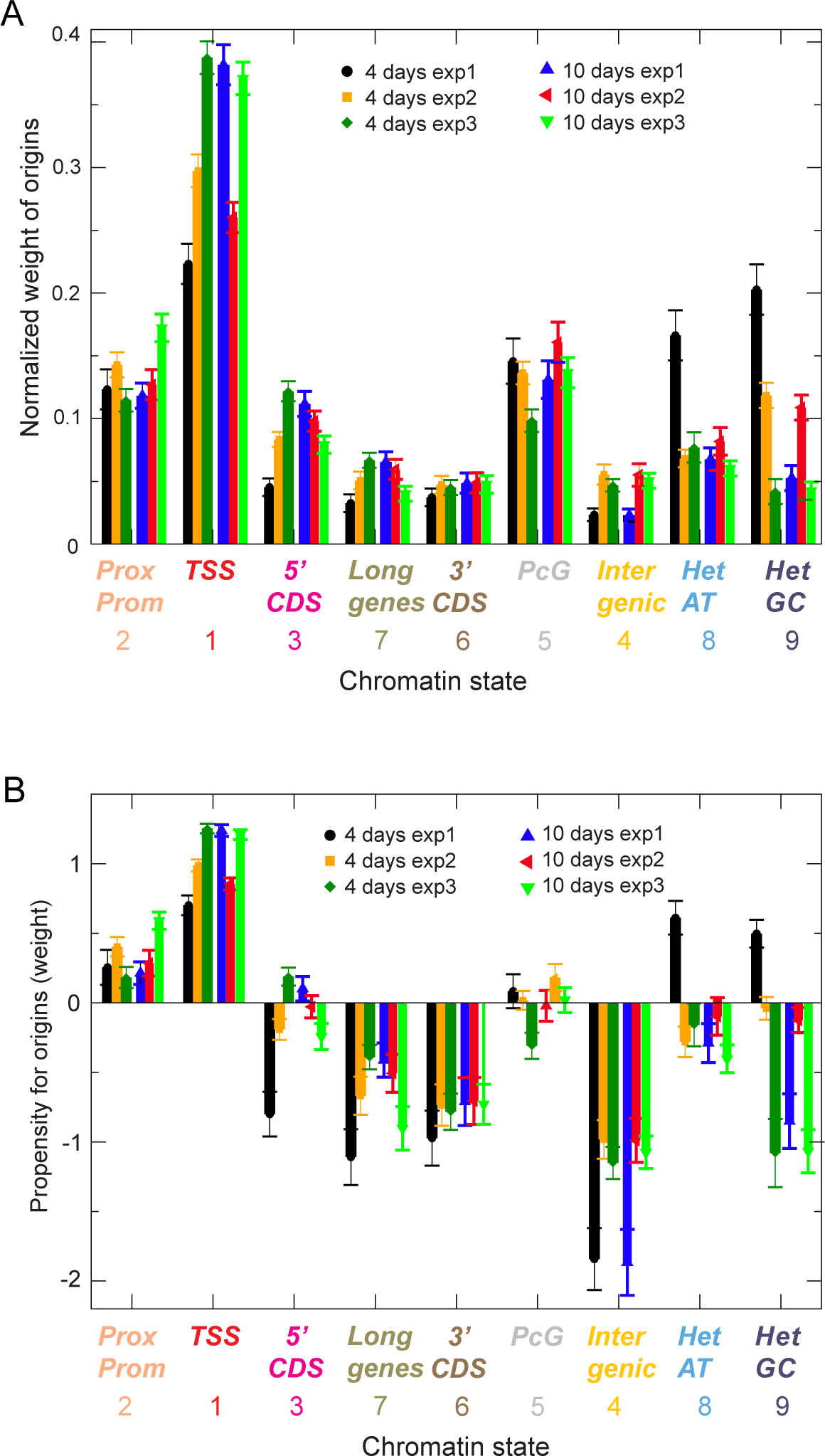
Association of ORIs with chromatin states. A. Normalized weight of ORIs belonging to the 9 chromatin states in each of the three independent experiments of 4 day-old and 10 day-old seedlings, as indicated. B. Same as in panel S4A, showing the propensity (instead of the cumulative weight) for ORIs in the 9 chromatin states is depicted for each of the three independent experiments of 4 day-old and 10 day-old seedlings, as indicated.

**Figure S5.**
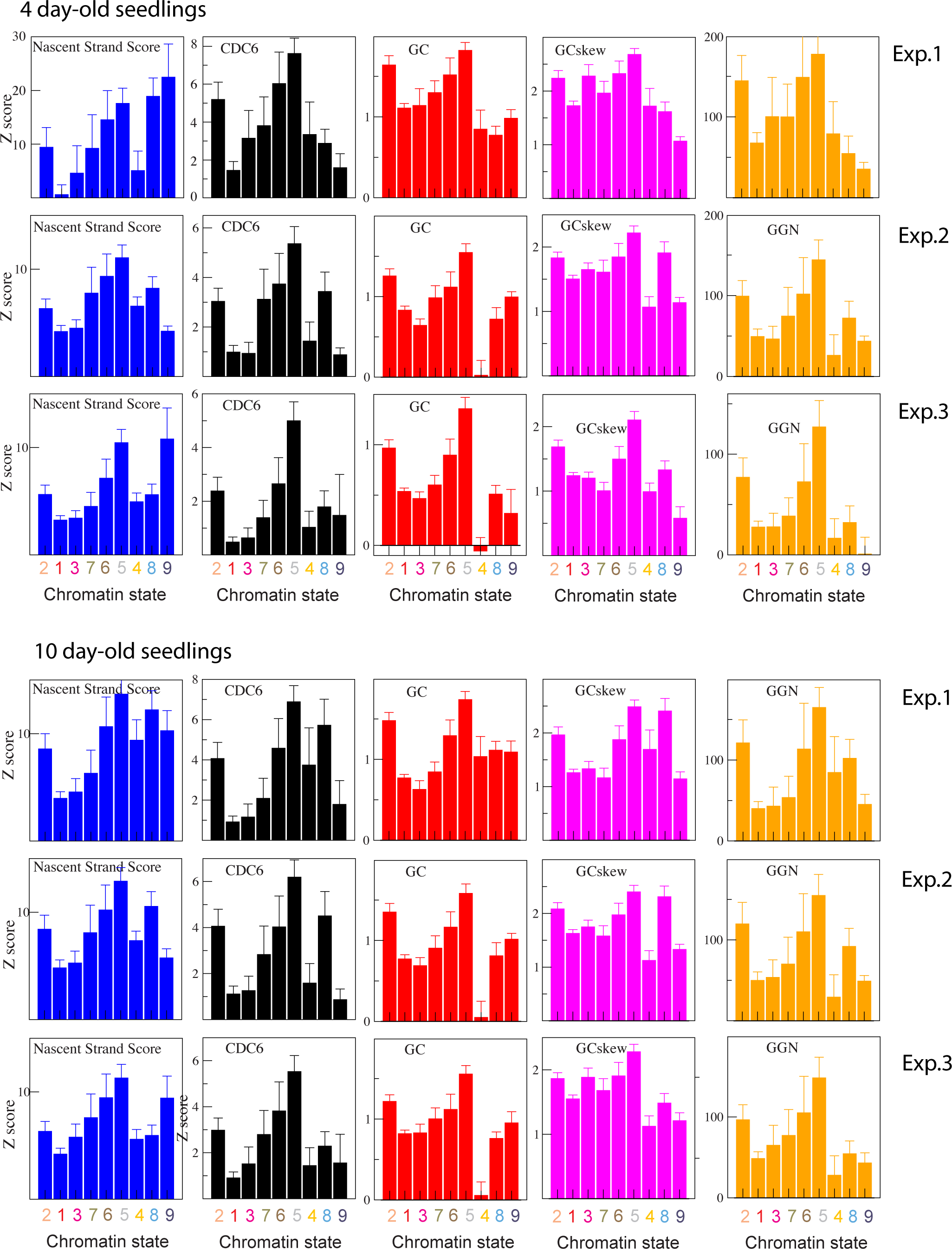
Relevance of several genomic variables for ORI specification. The weighted averages of the Z scores for several variables (NSS, CDC6, GC content, GC skew and GGN trinucleotide) are shown for each of the three independent experiments in 4 day-old and 10 day-old seedlings, as indicated.

**Figure S6.**
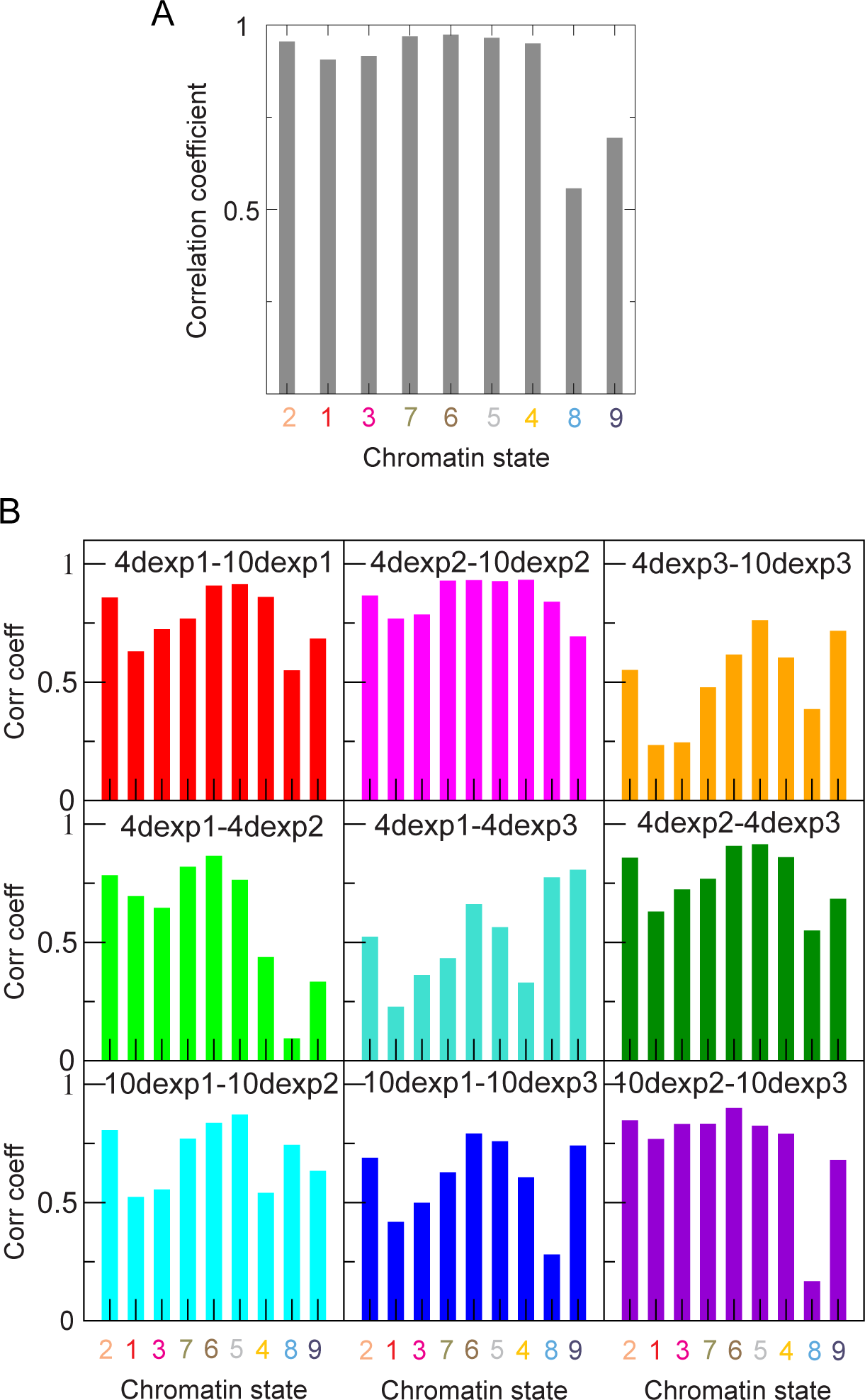
A. Correlation coefficients of the combined NSS of all ORIs identified in 4 day-old and 10 day-old seedlings, in each chromatin state. B. The correlations between the 6 individual NSS for the indicated experimental pairs.

**Figure S7.**
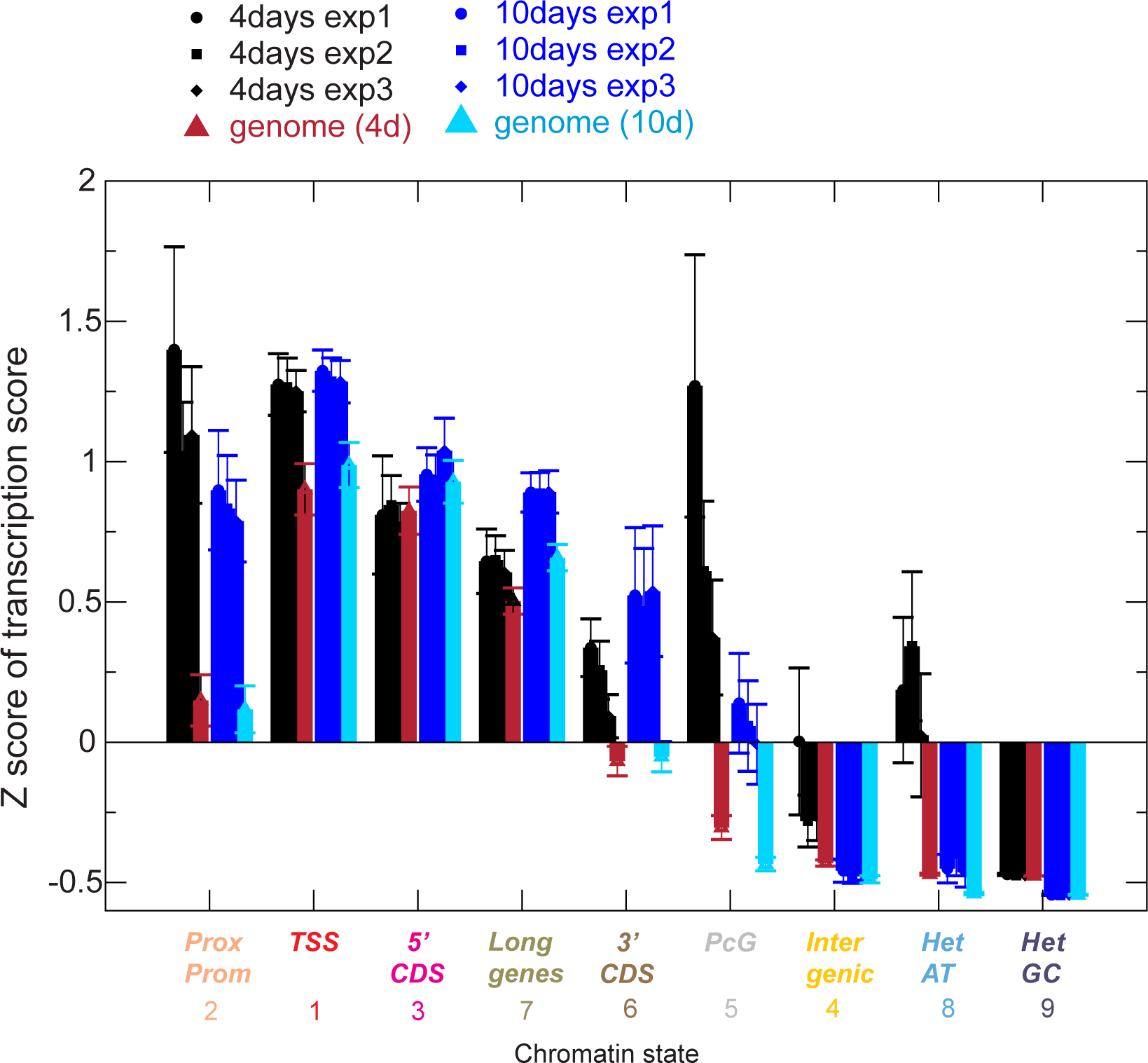
Average Z score of the transcription score with respect to the average transcription score of the entire genome is shown for ORIs in each chromatin state for each of the three independent experiments in 4 day-old (top panel) and 10 day-old seedlings (bottom panel), as indicated.

**Figure S8.**
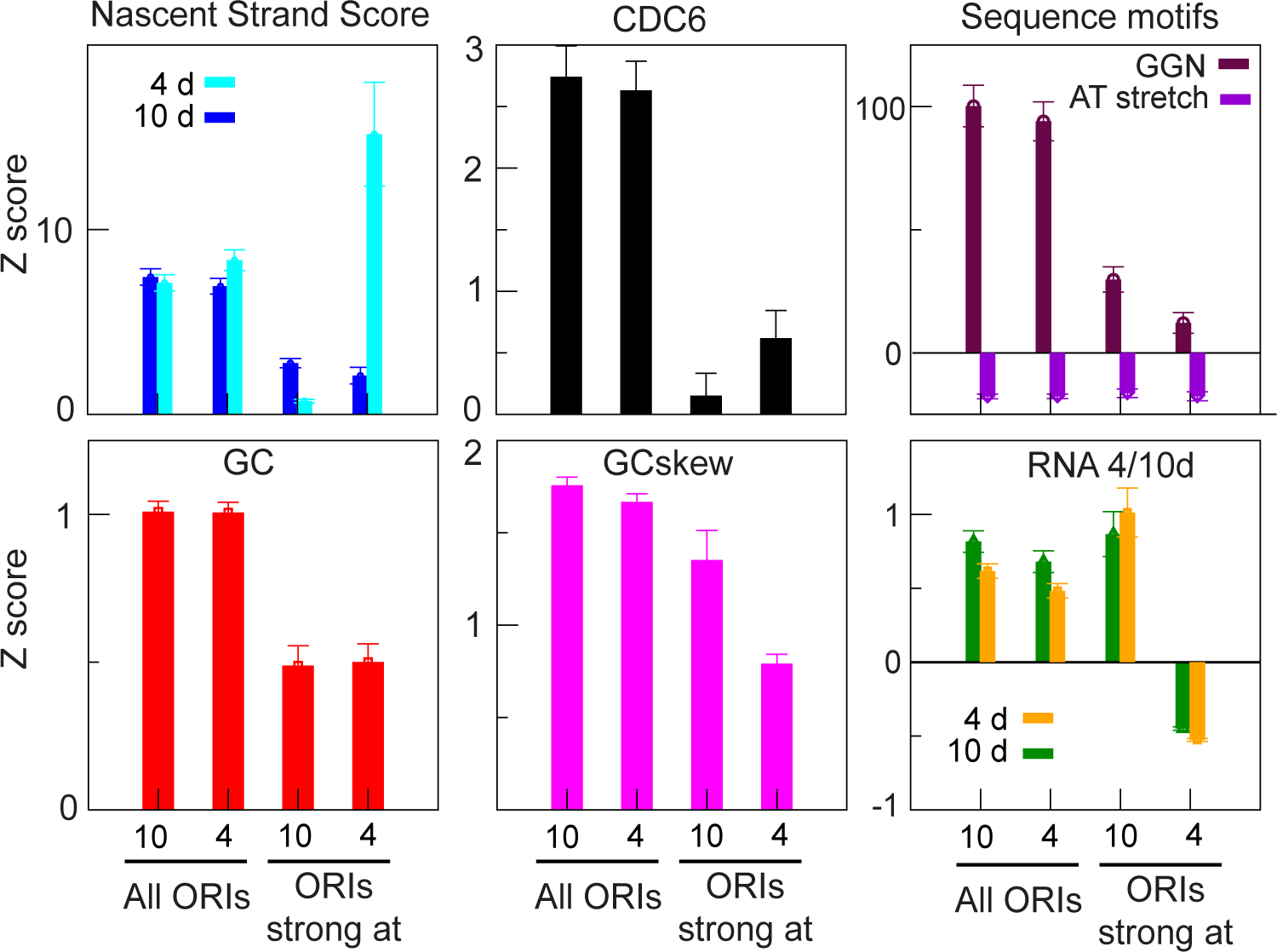
Weighted average of several genomic and epigenomic properties, using as weights the combined NSS, for the set of all ORIs and the ORIs preferred at 4 or 10 day-old seedlings, as indicated. In all cases we have used the stronger weight. Measurements were transformed to Z-score with respect to the whole genome, which produces positive values when they are higher than a generic genomic region.

### Supplemental Tables

**Supplemental Table 1. Genome localization of Arabidopsis ORIs.** The Table contains the chromosomal location, individual NSS values of each of the 2374 ORIs identified in each of the three independent experiments carried out in 4 day-old and 10 day-old seedlings, as well as the combined NSS of each developmental stage. Also the chromatin state associated with each ORI, according to the classes defined in Sequeira-Mendes et al. (2014) are also provided.

**Supplemental Table 2. Primers used to measure nascent strand abundance by qPCR.**

**Supplemental Table 3. Number and sequence of GGN motif in ORIs.**

**Supplemental Table 4. Developmentally regulated ORIs.** Genome localization of Arabidopsis ORIs preferentially active in 4 day-old and in 10 day-old seedlings with indication of their NSS values and the chromatin state associated.

